# LocAlign: Local Protein Structural Alignment with Geometric Deep Learning

**DOI:** 10.64898/2026.01.24.701262

**Authors:** Hagai Ravid, Jérôme Tubiana, Haim J. Wolfson

## Abstract

Identifying common function-determining structural motifs among proteins with different folds is a foundational task in computational biology with no go-to solution. Indeed, standard alignment tools like TM-align are ill-suited for matching small, sequence-order-independent motifs, while specialized tools have limited success. Here, we introduce LocAlign, a local structural alignment algorithm based on geometric deep learning. Given two protein structures, LocAlign iteratively predicts atom-level correspondences and a 3D superimposition. By formulating training as a weakly supervised task on pairs of proteins bound to identical ligands, we bypass the need for ground-truth alignments. We find that LocAlign recovers known functional motifs without explicit supervision, identifying high-quality alignments for 87% and 37% of protein pairs with similar and dissimilar folds, respectively. We show that, equipped with confidence scoring and motif-conditioning capabilities, LocAlign supports diverse applications, including functional annotation of the dark proteome and drug off-target screening. LocAlign is thus a potent, versatile framework for protein functional site comparison.

## 1 Introduction

Two evolutionarily unrelated proteins with a distinct three-dimensional fold may share similar molecular activity, such as binding to the same ligand or catalyzing a similar chemical reaction. This functional convergence is borne by local structural motifs, *i.e*., a handful of amino acid fragments with a conserved spatiochemical arrangement that mediate molecular activity. Classical examples of structural motifs include Cys2-His2 Zinc fingers, Calcium binding EF hands, catalytic triad of serine proteases, HTH motifs (nucleic acid binding), etc [Kessel and Ben-Tal, 2018]. Local structural alignment algorithms aim to detect such shared local structural motifs (or the absence thereof) between dissimilar proteins (Fig. 1). These algorithms are invaluable in basic molecular biology research for predicting shared (potentially unknown) molecular activity and pinpointing the corresponding functional motifs. The function-determining motif may later be used as a conditioning for generative design of functional proteins, using tools such as RFDiffusion [Watson et al., 2023, Butcher et al., 2025] or Chroma [Ingraham et al., 2023]. In drug discovery, local alignment algorithms can be used to compare ligand-binding pockets for drug repurposing and to predict off-target drug binding profiles [Yeturu and Chandra, 2011, Ehrt et al., 2016].

**Figure 1.**
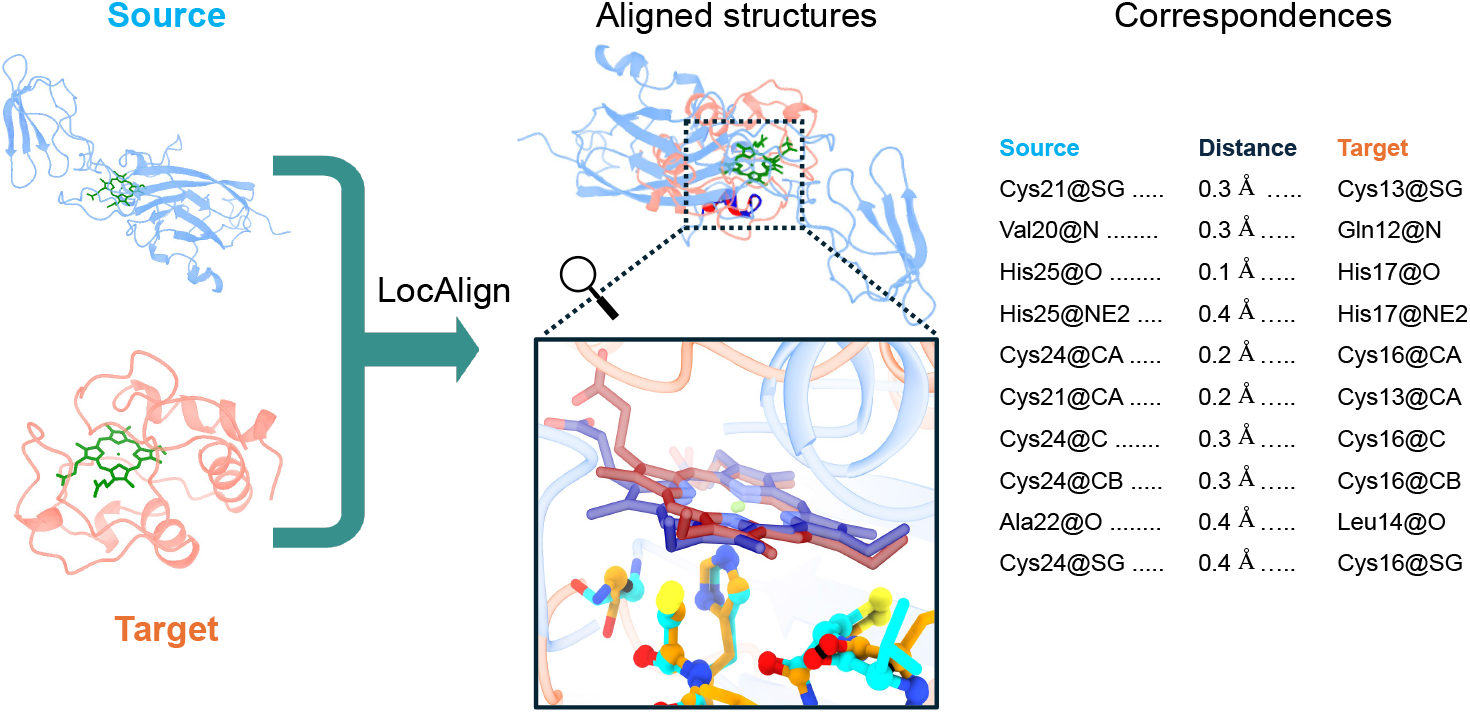
The local structural alignment problem. A local structural alignment between two proteins is a set of atom-level correspondences and a rigid-body transformation that superimposes them; high-overlap alignments suggest a conserved function and highlight the corresponding functional sites. The proteins are aligned in their *unbound* form, which means that the ligand is not part of the input. Shown here is an alignment of two heme C-binding proteins with LocAlign (PDBs: 1CI3:M, 1CO6:A; both belong to the validation set).

**Existing methods** can be categorized as follows. **Standard order-dependent structural alignment algorithms** such as TM-Align [Zhang and Skolnick, 2005] or FoldSeek [Van Kempen et al., 2024] are ill suited for this task because they assume sequence order dependence and thus can only detect domain-like structural motifs [Eidhammer et al., 2004]. DALI [Holm and Sander, 1995, Holm, 2022] and APoc Gao and Skolnick [2013a] relax the global sequence-order constraint, but still assume that matching linear fragments can be found. In contrast, US-align [Zhang and Pyle, 2022] provides a fully non-sequential (fNS) mode that treats proteins as 3D point clouds and employs a greedy algorithm to identify geometrically consistent patterns regardless of their sequential connectivity.

**Geometric hashing-based motif search tools** take as input a predefined local structural motif as template, and search for similar motifs in a structure database. The currently predominant algorithmic approach is based on Geometric Hashing [Lamdan and Wolfson, 1988] and dates back to the 90’s [Nussinov and Wolfson, 1991], with early implementations including TESS [Wallace et al., 1997], SiteEngine [Shulman-Peleg et al., 2004] and LabelHash [Pennec and Ayache, 1998], and modern works including the RCSB’s motif search tool [Bittrich et al., 2020] and the recently released FoldDisco [Kim et al., 2025]. While these algorithms excel at rapidly finding exact motif matches, they have several limitations. First, the functional motif must almost always be predefined in advance and thus, these algorithms are not suitable for protein-protein comparison, that is, motif discovery. Second, the motif should be relatively small, e.g., up to 10 and 32 residues for RCSB’s motif search tool and FoldDisco, respectively. Indeed, the number of computed geometrical features increases rapidly with the motif size (about 30-40 times the number of residues for FoldDisco), and in turn, so does the runtime and the expected number of false positives *E* (*E* ∝ *mn*, where *m, n* and the number of features in the motif and query database, respectively [Karlin and Altschul, 1990]). Third, the motif is assumed to be highly conserved in terms of amino acid types and relative orientations; allowed substitutions must be carefully defined to prevent overflow of insignificant hits. However, it is the rule rather than the exception that binding sites are extended and highly polymorphic, with conservation occurring at the chemical level (charges, hydrogen bond donors, etc.) rather than at the residue level.

### Deep learning-based protein alignment algorithms

There have been significant developments in deep learning for order-dependent sequence or structure alignments, e.g DEDAL [Llinares-López et al., 2023], BetaAlign [Dotan et al., 2023] and DeepBlast [Gonzalez et al., 2024]. However, order-independent alignment methods are rare, with SoftAlign [Trinquier et al., 2025] and the concurrent work PLASMA [Wang et al., 2025] being notable exceptions. In addition to the lack of reference architecture, a major obstacle is that there is only a limited number of gold-standard alignments of structural motifs. From a machine learning perspective, this amount of data is vastly insufficient to train a model in a supervised learning fashion. From a scientific perspective, this is an unsatisfactory state of affairs: in most cases of convergent function, there is no known function-determining local motif. It is plausible that in these cases, the motif is more complex and less conserved than in the “picture postcard” motifs aforementioned.

**Deep learning-based protein pocket comparison algorithms**, such as DeeplyTough [Simonovsky and Meyers, 2020] and PocketVec [Comajuncosa-Creus et al., 2024] also predict similarity between protein pockets using deep learning, but do not output structural alignments (atom-atom correspondences and 3D transformation). Thus, they lack interpretability and cannot be used in conjunction with motif-based generative protein design.

### Deep learning for 3D point cloud registration

At first glance, the local structural alignment problem can be seen as a special case of the *point cloud registration* problem in 3D Computer Vision, where atoms are viewed as colored points in space. Current state-of-the-art point cloud registration algorithms are all deep learning-based [Zhang et al., 2024]. Representative work includes Deep Closest Point [Wang and Solomon, 2019], Deep Best Buddies [Hezroni et al., 2021], RPMNet [Yew and Lee, 2020], GeoTransformer [Qin et al., 2022] and UDPReg [Mei et al., 2023]. These algorithms typically combine SE(3)-equivariant feature extraction, differentiable correspondence estimation, and rigid-body transformation solvers such as the differentiable SVD layer. In the protein domain, EquiBind [Stärk et al., 2022] applies similar principles to predict ligand–protein docking poses. While there are major differences between the two problems - most notably, the protein structures to be aligned are distinct objects with very low overlap - these considerations motivate us to pursue a deep learning approach to the local structural alignment problem.

Here, we introduce **LocAlign**, a geometric deep learning–based local structural alignment algorithm. Given two protein structures, LocAlign jointly predicts, in an iterative fashion, a correspondence between pairs of atoms and a rigid transformation that superimposes them in 3D. The model is end-to-end differentiable, SE(3)-equivariant and sequence order-independent, allowing it to detect conserved motifs even between evolutionary unrelated proteins. LocAlign supports both protein-protein and protein-motif comparison, and is fast enough for search against databases of moderate size (𝒪 (10^4^)). LocAlign is built on two main contributions. First, a reformulation of the local structural alignment problem as a scalable, weakly supervised learning task on pairs of proteins co-complexed with the same ligand; this bypasses the need for ground-truth alignments. Second, a custom deep learning architecture that combines building blocks of structural bioinformatics, 3D point cloud registration, and original modules. Our experiments show that LocAlign outperforms existing structural alignment tools, with the largest improvements observed on evolutionary unrelated protein pairs, where classical sequence-based or global structural alignment methods almost always fail. We show that LocAlign recovers well-known global and local alignments and uncovers many previously overlooked but visually convincing ones. Finally, we demonstrate through case studies that database search is feasible with LocAlign, and that it provides complementary information to traditional search with order-dependent alignment techniques.

## 2 Results

### 2.1 LocAlign: a Geometric Deep Learning Local Structural Alignment Algorithm

#### Model Architecture

LocAlign takes as input the sequence and *unbound* structure of two proteins (the source and the target), and optionally, per-atom motif masks. It outputs (1) a rigid-body transformation 𝒯 = (*R, t*) that superimposes the source’s ligand binding site onto the target’s binding site and (2) a list of weighted correspondences between the source and target atoms, 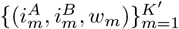, where the weights *w*_*m*_ are non-negative and sum to 1 and the number of correspondences *K*^*′*^ = 400 is formally fixed, but some can have zero weight. First, per-heavy atom embeddings are computed using two pretrained models: ESM2 [Lin et al., 2022], a sequence-based protein language model that captures sequence and structural context, and ScanNet [Tubiana et al., 2022], a geometric deep learning model for structure-based protein binding site prediction. These embeddings are concatenated and refined with a multi-layer perceptron (MLP). Next, a differentiable keypoint selection module, adapted from graph coarsening layers [Cangea et al., 2018], identifies keypoints atoms, typically within or around ligand binding sites, that are likely to be part of the alignment. This step reduces the computational complexity of the algorithm and forces the model to ignore irrelevant regions. Then, an initial soft correspondence matrix between keypoint atoms is computed. The matrix is such that two atoms have a high correspondence score - and thus should be closely superimposed in the *3D* space - if they are mutual nearest neighbors in the *embedding* space. The top-scoring correspondences are extracted, and geometrical inconsistencies between them are resolved by a Correspondence Solver Module (CSM). The CSM is a Graph Neural Network (GNN) where each correspondence is a node and edges encode geometrical consistency; it takes as input and outputs a per-correspondence importance weight. Next, the transformation that optimally super-imposes corresponding source and target keypoints is computed using a weighted Kabsch algorithm [Kabsch, 1976]. Finally, the transformation and correspondences are iteratively refined by a recycling mechanism [Jumper et al., 2021], that is, by feeding them back as extra inputs to the network.

Here, per-atom importance scores and positional encoding vectors are computed from the alignment and used to bias the keypoint selection and initial correspondence matrix steps, respectively.

### Training Data

Unlike in the sequence order-dependent case, there are very few ground-truth order-independent structural alignments to learn from - mostly classical motifs such as classical catalytic triads or metal coordination sites. On the other end, there is plenty of protein pairs for which a high-quality local alignment *presumably* exists: proteins that bind the same ligand. We reasoned that the 3D transformation that superimposes the corresponding structural motifs should also, to some extent, superimpose the bound ligands with low root mean square deviation (Fig. 1). Given a pair of *unbound* protein chain, our supervised learning task will consist in finding a transformation that best superimposes the two *unseen* ligands. Using the Protein Data Bank and the BioLiP database [Zhang et al., 2023], we compiled a non-redundant and balanced dataset of about 120.000 pairs of protein chains, covering over 800 distinct ligands.

#### Training loss

The model is trained using two complementary objectives: a supervised **ligand loss** and an unsupervised **alignment quality loss**. The ligand loss ℒ_lig_ is the Root Mean Square Deviation (RMSD) between the poses of the target ligand and of the source ligand, after applying the output transformation. It incentivizes the model to find motifs that are conserved from the point of view of the ligand, and thus likely to be essential for binding. The alignment quality loss, which combines three terms, incentivizes the model to find alignments that are interpretable and biologically significant. The first term, ℒ_corr_, is the (weighted) RMSD between pairs of atoms in correspondence; it is such that corresponding atoms should be close to each other in 3D space, after the transformation is applied. The second term, ℒ_emb_, is a contrastive loss adapted from CLIP [Radford et al., 2021]; it is such that corresponding atoms should be close to each other in the embedding space. ℒ_emb_ encourages correspondences between atoms of similar chemical type and local environment. It also serves as a representation learning loss: atoms that are in correspondence (*e.g*., because they are closely-superimposed in 3D) should also have high embedding similarity. The third term, ℒ_gyr_ is the scaled radius of gyration of the set of aligned atoms; it promotes more compact alignments. The total loss is given by:

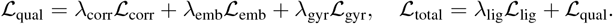

The learning task is weakly supervised in the sense that there is only supervision on the transformation and not on the correspondences. Moreover, the transformation loss does not necessarily have a single global minimum: for single-atom ligands there are infinitely-many transformations that superimpose them with zero RMSD and for flexible ligands with hinges, there could be two distinct transformations with similar RMSD. Detailed descriptions of the architecture and training procedures are provided in **Methods**.

### 2.2 LocAlign recovers global and local order-dependent alignments

We first showcase examples of high-quality alignments inferred by LocAlign.LocAlign successfully recovers classical global and local *order-dependent* alignments, even though there is no order-dependent constraint and no *explicit* supervision. LocAlign accurately superimposes Human Ubiquitin to Human SUMO1 - a classical example of proteins with high structural but very low sequence identity 16% [Van Kempen et al., 2024] (Fig. 3a). Accordingly, the local alignment focuses on the conserved beta-sheet, and mostly involves backbone atoms (64/75 of the top correspondences). The model also recovers small domain-domain alignments, such as one between two Cys2-His2 zinc fingers of unrelated proteins (Fig. 3b). The alignment is centered around the zinc ion, involves both backbone and side chain atoms and has sub-angstrom resolution (ligand RMSD: 1.2 Å, correspondences RMSD: 0.9 Å). We stress that ligands are not part of the input, and hence, the fact that the model accurately superimposes them from the structure information alone is non-trivial. Close-up inspection reveals that the two histidines and two cysteines that chelate the zinc ion are closely superimposed and in correspondence. Two of the histidine pairs and one of the cysteine pairs have multiple atoms among the top correspondences, while the last cysteine pair, which has higher RMSD and lower correspondence weight (rank: 112), is not shown.

**Figure 2.**
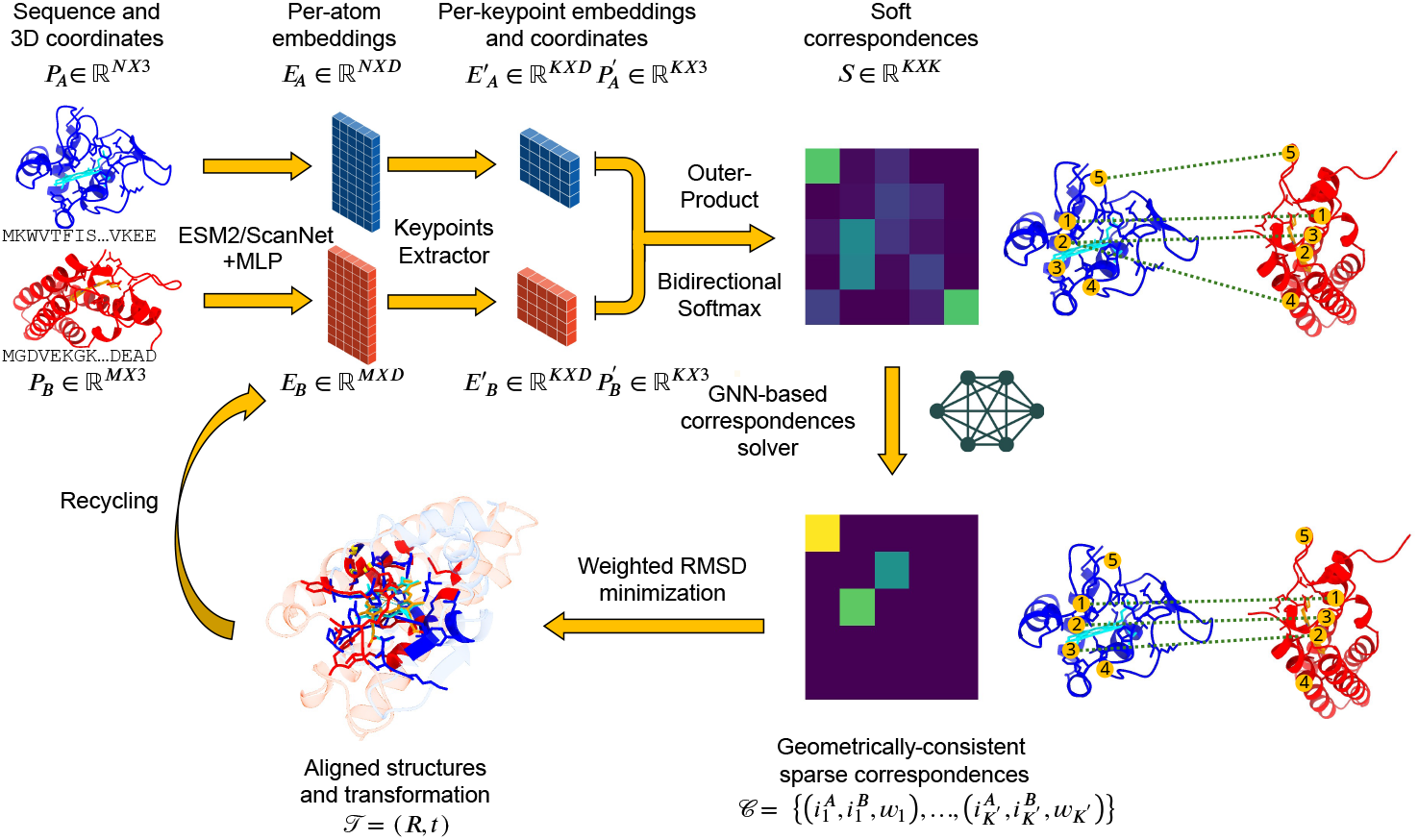
Model Overview: Given the sequence and structure of two proteins, LocAlign first extracts peratom embeddings using pretrained models and refines with an Multi-Layer Perceptron (MLP) block. Next, a differentiable keypoint selection module identifies keypoint atoms that are likely to be part of the alignment. A soft keypoint-to-keypoint correspondence matrix is calculated and subsequently sparsified and refined. The correspondences are fed to a differentiable, weighted Kabsch algorithm that finds the rigid transformation that best superimposes keypoint pairs. Finally, a recycling mechanism allows to iteratively refine the transformation. The ligands are not processed by the model and shown only for illustrative purpose.

**Figure 3.**
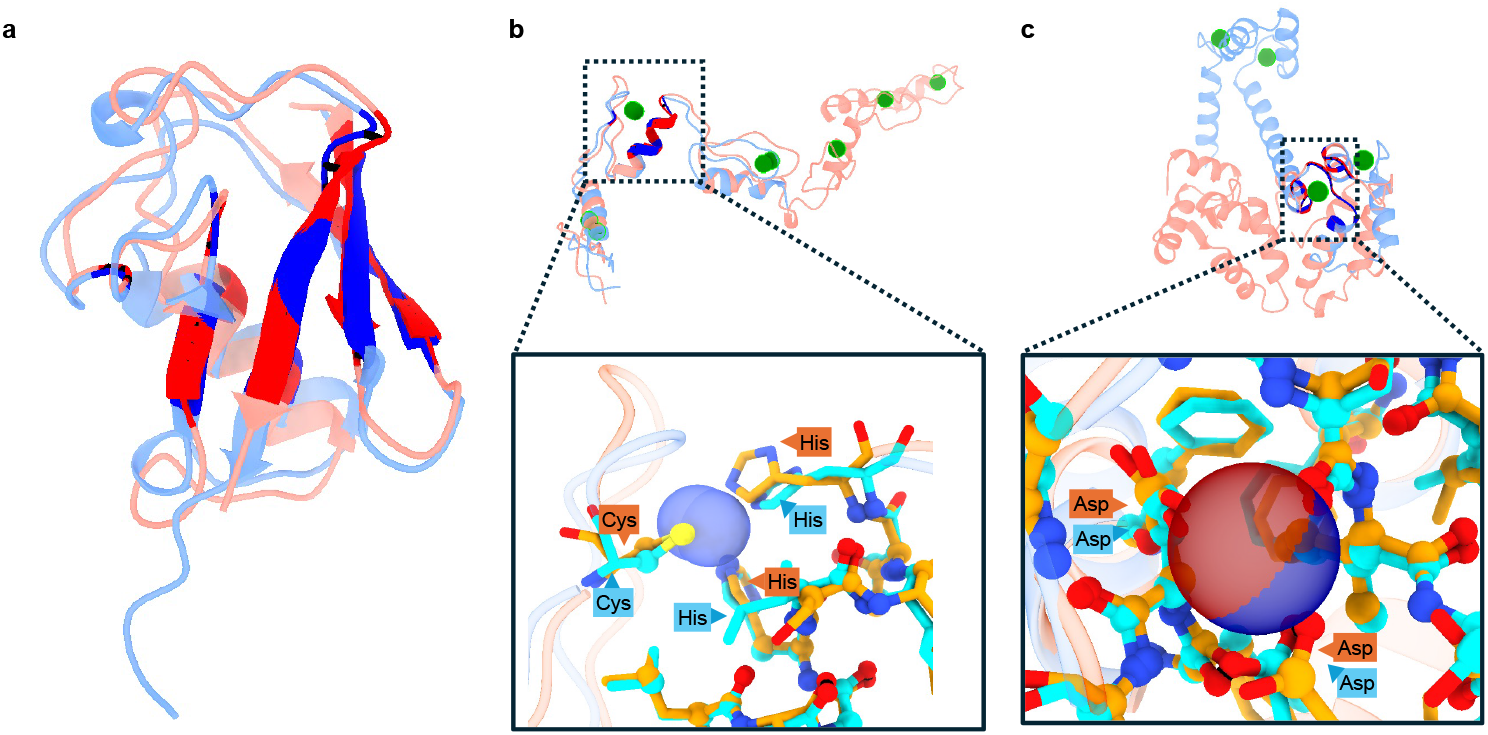
Examples of alignments recovered by LocAlign. (a) Human Ubiquitin and SUMO1 (PDB IDs: 1ubq:A, 1wm3:A); (b) Zinc-bound EGR1 and TFIIIA (PDB IDs: 1a1F:A, 1tf6:A); (c) Calcium-bound Human Calmodulin and Recoverin (PDB IDs: 1cll:A, 2het:A). In each example, the target and source proteins are shown in red and blue color palettes, respectively. For (a), the alignment is shown in a single global view. For (b) and (c), the top inset panel shows a global view with structures in cartoon representation, ligands in green sticks, and aligned residues highlighted; the lower panel focuses on the aligned motif, where aligned residues are shown in opaque sticks and others in transparent cartoon. Within the aligned residues, the actual atoms that are in correspondence are shown as balls, with black links indicating the mapping. For visual clarity, only correspondences with the largest weights are depicted, using an adaptive inclusion threshold such that 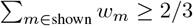.

LocAlign can also successfully align short linear motifs within proteins of different folds, such as the Cytochrome C heme binding motif (CXXCH, Fig. 1) and an EF-hand motif (Fig. 3Cc. Aligning Human Calmodulin and Recoverin, two flexible calcium-binding proteins, reveals a highly conserved EF-hand motif (ligand RMSD: 0.15 Å, correspondences RMSD: 0.64 Å), in which the Calcium ion is tightly chelated by a network of Aspartic acid side chains, backbone carboxyl groups and amide groups. More examples of order-dependent alignments are shown in Supplementary Figure 3.

### 2.3 LocAlign uncovers novel order-independent alignments

We next present examples of order-independent alignments, where the two proteins do not have the same fold. Fig. 4a depicts a high-quality alignment of two iron sulfur cluster chelation sites. In both cases, the chelation site is conformational, that is, constituted by residues from two distinct linear fragments, and the relative orientation of the two fragments is not conserved. Nevertheless, the model identifies a shared chelation motif of two cysteines and one alanine, as well as an additional correspondence between a cysteine and histidine side chain. Indeed, histidine and cysteine are often interchangeable in ion chelation site, making this correspondence plausible. We found several other examples of ion chelation sites with substitutions “accepted” by the model (Supplementary Fig. S4).

**Figure 4.**
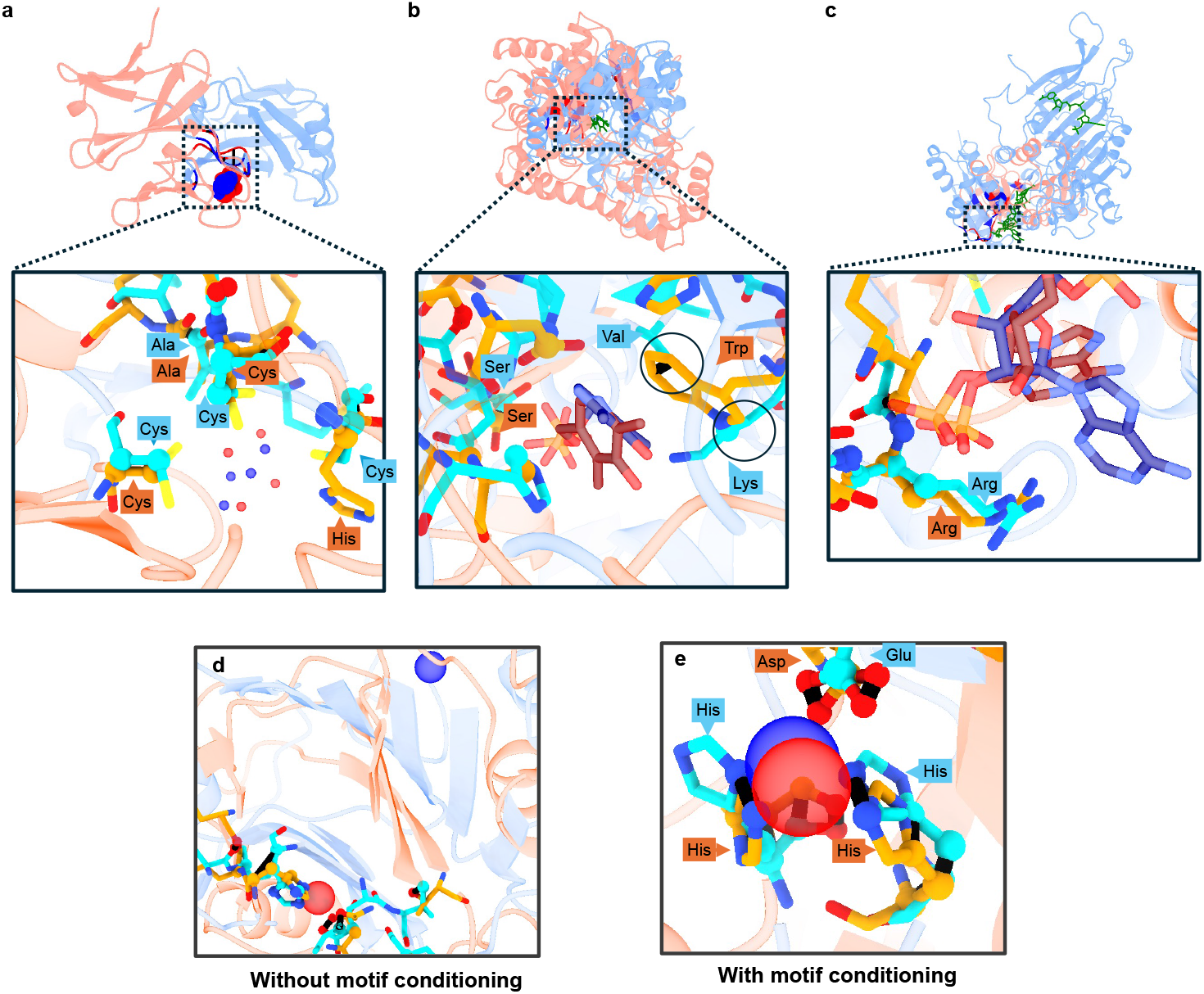
LocAlign identifies order-independent alignments and chemical substitutions across diverse folds. (a) Alignment of iron-sulfur cluster (FES) chelation sites (PDB IDs: 1x0g:C, 3gce:A). (b) Vitamin B6 Phosphate (PLP) binding pockets showing a two-to-one substitution (PDB IDs: 1rcq:A, 7aat:A). (c) NADP binding pockets (PDB IDs: 2b9h:A, 5sda:A) showing atomic-level correspondence in a flexible ligand context. (d,e) Alignment of iron (Fe^2+^) chelation site, with (d) and without (e) using motif conditioning (PDB IDs: 1ey2:A, 5j92:B). Providing the source’s chelating residues as motif guides the model towards the desired alignment.

Fig. 4b presents an alignment of two Vitamin B6 Phosphate binding pockets. Although the transformation found by the model closely superimposes the two ligands, the respective pockets are much less conserved. Except for a conserved serine forming hydrogen bonds with the phosphate group, most correspondences involve closely superimposed atoms from residues of different types. Interestingly, the pockets feature a *two-to-one substitution*, where a large tryptophan side-chain in one pocket is substituted by a valine and a lysine side chains in the second. This case exemplifies the benefit of atomic-resolution alignments, where the geometry and chemical properties of the pocket are conserved, but not its amino acid content. We found other examples of two-to-one examples, such as a copper ion chelation site in which two histidine side chains are substituted by the backbone amide and side chain of a single histidine (Supplementary Fig. S4c).

Finally, we report in Fig. 4c an alignment of two pockets binding Nicotinamide-Adenine-Dinucleotide Phosphate (NADP) - a large and flexible ligand. In this case, no rigid transformation can perfectly superimpose the ligands and, accordingly, the model only aligns well the central part. A conserved arginine at the entrance of the pocket is identified, together with many atomic-level correspondences that delineate the pocket.

### 2.4 Local supports motif-conditioned alignments

In many workflows, we seek to align user-defined motifs instead of full proteins. Classical algorithm address this use-case by discarding all non-motif atoms from the input, loosing valuable contextual information in the process. In contrast, LocAlign preserves the complete structure to compute context-enriched embeddings, and only afterwards uses the motif to guide the keypoint selection. We found that in many instances, motif conditioning could rescue bad or undesired alignments. One such case - a pair of Iron-bound proteins - is reported in Fig. 4(d,e). Without conditioning, LocAlign aligns a beta sheet with high geometric similarity but no functional relevance. Using one of the Iron chelation sites as motif, LocAlign perfectly aligns the two *Fe*^2+^ sites, revealing conserved histidines and a benign glutamate to aspartate substitution.

### 2.5 Quantitative performance evaluation

We next benchmarked LocAlign against three classical structural alignment tools (TM-Align [Zhang and Skolnick, 2005], DALI [Holm, 2022], and APOC [Gao and Skolnick, 2013a]), one modern implementation (US-Align-2 [Zhang and Pyle, 2022]), and two deep learning-based approaches (SoftAlign [Trinquier et al., 2025] and PLASMA [Wang et al., 2025]). Geometric hashing-based methods (FoldDisco, RCSB’s motif search tool and SiteEngine) were not included here due to size limitations: for the first two, an input motif of size at most 32 or 10 residues must be provided, and for the last one, the lengths of the target and source proteins were limited to *L*_tar_ ×*L*_source_ *<* 50.000.

#### Success Definition

An alignment prediction is considered successful if it meets three simultaneous criteria: (1) The root-mean square deviation (RMSD) between the poses of the target ligand and the *transformed* source ligand pose is below 4Å ; (2) the (weighted) RMSD between corresponding atoms is below 2Å, that is, the motifs are tightly superimposed. (3) Corresponding atoms are significantly more often of the same heavy atom type than expected for a random pairing (Methods). Examples of unsuccessful alignments are shown in Supplementary Figs. S2,S5,S6.

#### Data Stratification

The difficulty of the alignment task depends on the degree of similarity between the target and the source protein. To quantify it, we used the CATH hierarchical classification system [Waman et al., 2024]: the two proteins were considered of the same fold if at least one of their respective domains belonged to same homologous superfamily (H-level). We also investigated finer-grained stratification based on the other levels of the CATH hierarchy, but found that the difficulty mostly depended on the H-level. Distributions of evaluation metrics on the fine-grained stratification are shown in Supplementary Fig. S7.

#### Data Partition

The difficulty of the *learning* task depends on how similar are the test examples to the training examples. We evaluated the performance under two complementary data splits. **Homology split:** Proteins were clustered using MMseqs2 [Steinegger and Söding, 2017] with a threshold of 50% sequence identity and 60% coverage. Pairs assigned to the test set share no clusters with the training set, ensuring that the model must generalize beyond homology. **Ligand split:** None of the ligands in the test set appeared in the training set. This is a much harder generalization task, since the model must align protein binding sites without having ever seen their shared ligand.

Table 1 summarizes success rates for the two data splits, stratified by fold similarity. For proteins with similar folds, LocAlign, TM-Align, US-align and DALI all performed relatively well, with success rates above 85% and 70% for the ligand and homology splits. The fact that LocAlign was better or on par with these methods is remarkable, as the order-dependent constraint is generally critical for accurate alignment of structures [Trinquier et al., 2025]. Conversely, SoftAlign and PLASMA had lower success rate for this task, presumably because they do not explicitly estimate the correspondences and the transformation (i.e., they lack the rigid superimposition constraint). APoc performed poorly in our experiments, presumably because of failures at the pocket extraction stage. Visual inspection of unsuccessful LocAlign alignments mostly revealed benign failure modes, such as miscalculated ligand RMSD for symmetric and multivalent ligands, or aligning the non ligand-binding domain of multi-domain proteins (Supplementary Fig. S5).

**Table 1.** Success rates (%) for baselines, LocAlign, and ablations. For LocAlign, success is defined as ligand RMSD ≤ 4 Å, atom type acceptance *>* 0.5, and correspondence RMSD ≤ 2 Å . ^*†*^For the baselines, success is defined by ligand RMSD ≤ 4 Å only; these methods were not filtered for correspondence RMSD. Bold indicates best performance; underlined indicates second-best.

| Group | Method | Homology split |  | Ligand split |  |
| --- | --- | --- | --- | --- | --- |
| | | ≠ fold | same fold | ≠ fold | same fold |
| Baselines<sup>†</sup> | DALI | $\leq 2.2\%$ | 71.9% | $\leq 1.5\%$ | 85.9% |
| | TMalign | $\leq 2.3\%$ | 70.5% | $\leq 1.4\%$ | 86.3% |
| | APoc | $\leq 0.8\%$ | 7.1% | $\leq 0.4\%$ | 8.6% |
| | US-align_sns | $\leq 2.7\%$ | 73.3% | $\leq 1.5\%$ | 86.3% |
| | SoftAlign | $\leq 0.4\%$ | 30.2% | $\leq 0.8\%$ | 63.3% |
| | PLASMA | $\leq 0.3\%$ | 19.6% | $\leq 0.5\%$ | 40.0% |
|  | LocAlign (ours) | **37.0%** | **87.0%** | <u>13.1%</u> | <u>87.0%</u> |
| Ablations | w/o ESM embeddings | 13.1% | 81.5% | 4.4% | 85.9% |
|  | w/o GNN | <u>35.4%</u> | **87.0%** | **13.7%** | 84.9% |
|  | w/o ligand loss | 2.0% | 54.6% | 0.9% | 53.8% |
|  | w/o quality loss | 1.3% | 50.7% | 0.4% | 57.9% |
|  | w/o recycling | 6.2% | <u>86.5%</u> | 3.5% | **88.0%** |

For dissimilar folds, the difference is dramatic: all baselines almost entirely failed, whereas LocAlign reached 37% success rate on the Homology split. The success rate drops to 13% on the ligand split, indicating that generalization to unseen ligands is much harder, but not impossible. This drop echoes recent evaluations of AI-based protein-ligand docking algorithms [Buttenschoen et al., 2024, Jain et al., 2024, Oruç et al., 2025]. The lower success rates reflect the difficulty of the local, order-independent structural alignment task. First, the search space is much larger, as there is no order-dependent constraint and one-to-one mapping at the amino-acid level. Second, some binding sites are poorly conserved (e.g., due to different binding modes, or to the ligand being flexible) and simply cannot be well aligned, as previously shown for ligand binding pockets [Gao and Skolnick, 2013a,b]. We next performed ablation studies of the various building blocks of LocAlign. Removing either the supervised ligand loss or the unsupervised alignment quality loss resulted in a large drop in success rates. Visual inspection revealed that without ligand loss, the model prefers to align common structural motifs such as helixes and beta sheets rather than ligand binding sites (Supplementary Fig. S2 B). Conversely, without the alignment quality loss, the model correctly superimposes the ligands, but the correspondences are underdetermined and of low quality (Supplementary Fig. S2 C). Regarding the architecture, removing the ESM2 embeddings (leaving only ScanNet embeddings) led to a sharp decrease in performance, indicating that high-quality initial embeddings are essential, and that perhaps end-to-end fine-tuning could further boost performance. The recycling mechanism was determinant to achieve high-quality alignments in examples with different folds, and the Correspondence Solver Module was also found to be beneficial overall.

### 2.6 Model Confidence

These results demonstrate that LocAlign can produce novel, high-quality alignments, while also failing in many cases. For practical usability, it is therefore imperative to also output a confidence score that can be used to filter low-quality alignments. By analogy with AlphaFold’s pLDDT and pAE heads, we propose using the predicted ligand RMSD (pLRMSD) as a confidence score. In our implementation, the pLRMSD is calculated by aggregating four alignment quality scores with a monotonic Generalized Additive Model (GAM), trained on the validation set. The pLRMSD is low (*i.e*., higher confidence) for alignments that have high embedding and geometric similarities, that are compact and of large size (Methods and Supplementary Fig. S8a). We found that pLRMSD correlated well with LRMSD (cross-validation Spearman’s correlation coefficient: *ρ* = 0.59, Supplementary Fig. S8b and that filtering alignments by pLRMSD was a valid strategy. Indeed, while only 37% of alignments were successful for pairs with different folds, the success rate increases to 80% (resp. 65%) after retaining alignments with pLRMSD ≤ 2Å (resp. 4 Å), (Supplementary Fig. S8c. Accordingly, more than 80% of pairs with similar folds had pLRMSD ≤ 2Å . Taken together, these findings suggest that LocAlign, paired with a model confidence head, can be of practical utility for day-to-day structural bioinformatics analysis.

### 2.7 Proof-of-concept applications to database search

We last investigated whether LocAlign would be practically useful in a *database search* setting, in which we compare a query structure (or motif) against a large database of template structures (or motifs). To limit the run-time, we first pre-compute initial per-atom embeddings for all templates and cache them for future database queries. For ranking alignments, we hypothesized that, much like AlphaFold’s high-pLDDT score is indicative of whether a protein is globular or disordered (i.e., it does not have a stable 3D structure), a low pLRMSD score should be indicative of whether or not the two proteins share a fold or a ligand binding sites.

We first constructed a non-redundant database of ∼ 33.000 templates from the BioLIP2 database and ran a few search queries against the database (see detailed Methods). The preprocessing was completed in 140 minutes, followed by 12-25 minutes per query on a Nvidia L40S GPU, depending on the size of the structure. The per-query runtime is thus reasonable, and much faster than the time it would take to run a virtual screening with AlphaFold3 or Boltz-2 against the BioLIP2 ligand set (roughly 10-30s per ligand). Regarding the top hits, as expected, we found that whenever the query itself was present in the database, it was ranked first or among the top hits according to the pLRMSD metric and whenever structural homologs existed, they were also hits. For instance, querying 9cte:A, a de novo design of porphyrin-containing enzyme [Hou et al., 2025] yielded the 9cte:A itself (pLRMSD=1.06, rank 1), the original porphyrin-binder from which the computational design was initiated (7jrq:A, pLRMSD=1.83, rank 3) [Mann et al., 2020] and other porphyrin binders (Supplementary Fig. S9a).

We next assessed whether LocAlign could be used to annotate ligand binding sites in more challenging cases where traditional structural comparison tools are of no help: the dark proteome. Barrio-Hernandez et al. [Barrio-Hernandez et al., 2023] recently used FoldSeek to systematically analyze the AlphaFold database, and found *>* 700K “dark clusters” - groups of proteins with no detectable structural similarity to any protein family with known functional annotations. For each query, we compare it against a database of *template motifs*, defined as the ligand binding sites of the non-redundant BioLIP2 database. Hits are ranked by the pLRMSD score, with a per-ligand rescaling to correct for ligand size bias (Methods). We queried representatives of 14 dark clusters - the four examples showcased in Figure 2 of [Barrio-Hernandez et al., 2023] and computationally annotated with DeepFRI [Gligorijević et al., 2021], and the ten examples showcased in Figures 4,5 of [Fang et al., 2025], annotated with ATOMICA. Five queries (A0A0M0BG38,A0A2D8BRH7,A0A849TG76,S0EUL8,V5BF69) returned low-confidence hits, with high pLRMSD or low-quality alignments (Supplementary Fig. S9b). Four of the five heme C binders identified by ATOMICA and experimentally-validated (A0A1T4N4K0,A0A2V6P8N7,A0A7W0X6V6,A0A7W1B5T5) had heme C binding-proteins as top hits (Supplementary Fig. S9c); inspection of the local alignments revealed a classic cytochrome C-binding motif (CXXCXXH). Similarly, the ATOMICA-predicted heme binder (Fig. 5a, A0A6B2YTG2, and three zinc binders (A0A6C0E0P4,A0A7Y4VG56, W8VVQ2, Fig. 5b) were also confirmed by LocAlign. For the zinc binders, a classical Glu2His2 chelation site was found. The last query, A0A849ZK06, yielded intriguing results (Fig. 5c). Although DeepFRI predicts a ribonucleotide binding activity, we could not corroborate this prediction using LocAlign or the nucleic acid binding site predictor PeSTO [Krapp et al., 2023]. Instead, we found alignments with metal chelation sites (Zinc, Manganese or Nickel, a prediction corroborated by ZincSight [Mechtinger et al., 2025]), and intriguingly, binding sites of three metalloproteinase inhibitors. Close-up inspection revealed that: (1) Several of the top hits are catalytic sites of metalloproteinases, including the Human ACE2 protein and (2) The putative metal ion was only partially coordinated by one glutamate, one histidine and one oxygen molecule (which is itself stabilized by a glutamate and an arginine), whereas the second histidine found in aligned sites was mutated to valine - a classical feature of catalytic metals. Our analysis thus suggests that A0A849ZK06 is a metalloproteinase with high local, but low global structural similarity to other known metalloproteinases.

**Figure 5.**
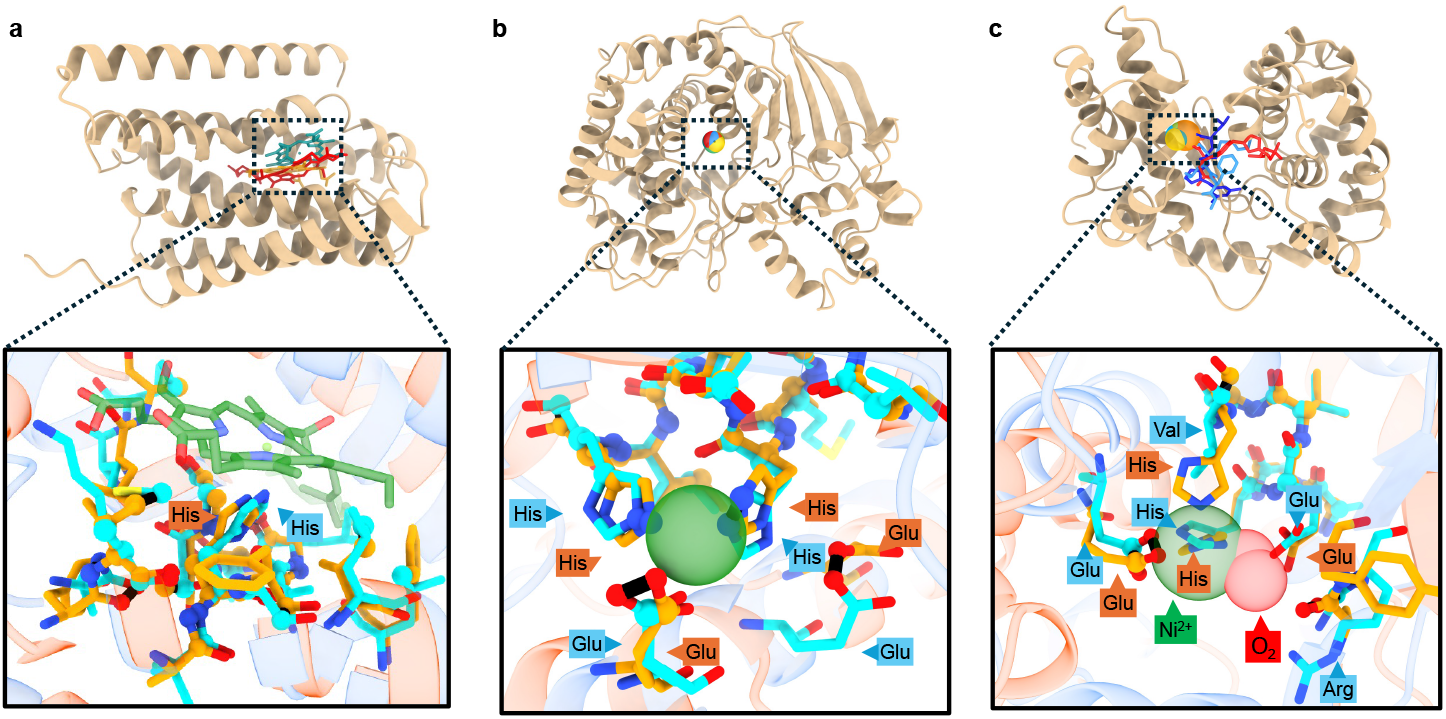
LocAlign applications to database search and the dark proteome. (a) **Heme binding prediction:** For the dark cluster representative A0A6B2YTG2, LocAlign identifies a high-confidence match to a Heme A binding site (template 7rjo:A, pLRMSD = 2.95Å). (b) **Zinc binding site identification:** Querying the dark cluster protein A0A6C0E0P4 yields a near-perfect alignment (pLRMSD = 0.00Å) to a known Zinc binding motif (template 5d7w:A). (c) **Functional reannotation of A0A849ZK06:** LocAlign identifies matches to multiple metalloprotein catalytic sites (shown here is template 5jig:A, pLRMSD = 0.52Å, ligands: OXY and NI).

Finally, we investigated whether LocAlign could be useful to compare drug binding pockets, for off-target binding prediction / repurposing. Imatinib is a FDA-approved anti-cancer drug that blocks the Bcr-Abl kinase, as well as other kinases. We queried the kinase domain of Bcr-Abl (1iep:A) using the imatinib binding pocket as motif, against the AlphaFold-predicted human proteome [Varadi et al., 2024] (23 586 templates, including fragments of long proteins). As shown in supplementary Table S1, We found that, among the top-40 hits, 19 are known interactors, including Bcr-Abl itself (rank 4/23,586), a known repurposing target (C-Kit, rank 31), and many known off-targets (e.g., DDR, PDGFR ranked 13,21/23,586). The results are thus qualitatively valuable, though, for optimal accuracy, LocAlign should be integrated together with AlphaFold3 or Boltz-2 [Passaro et al., 2025] in a multi-step virtual screening pipeline.

## 3 Discussion

The local structural alignment is a basic task in structural bioinformatics with currently no go-to solution. Classical order-dependent alignment algorithms such as TM-Align mostly only succeed when chains have similar fold, and thus fail to detect shared functional sites among evolutionary-unrelated proteins. Conversely, specialized order-independent methods have poor success rate due to the vastly larger search space. Additionally, even modern works such as FoldDisco [Kim et al., 2025] remain limited to residue-level representations, often overlooking the delicate atomic-scale patterns critical for function. Here, we proposed LocAlign, a first-in-class deep learning-based local structural alignment algorithm. Our main contributions are: (1) a differentiable architecture that is transformation-equivariant, sequence order-independent, atomic-scale, and computationally efficient, and (2) a weakly supervised learning protocol that bypasses the need for ground-truth alignments. LocAlign performs equivalently or better than standard algorithms on fold comparison tasks despite lacking the order-dependent prior, and, more importantly, can detect conserved structural motifs even when there is no global fold similarity. Equipped with motif-conditioning and model confidence prediction capabilities, LocAlign already proves to be a useful tool for day-to-day structural bioinformatics analysis. Using it, we notably uncovered previously unreported patterns, such as two-to-one residue substitutions within ligand binding or ion chelation sites, where chemistry is preserved at the atomic rather than the residue level. We anticipate that LocAlign and future versions could be valuable for a variety of tasks, including protein function annotation, drug repurposing, drug off-target binding prediction and motif-based generative protein design. An inherent limitation of our approach is the assumption that proteins that bind to the same ligand share similar binding sites that can be rigidly aligned. Although chemically plausible, this hypothesis does not always hold [Gao and Skolnick, 2013b], and thus the achievable success rate is upper-bounded. Nevertheless, it should be possible to improve the performance of LocAlign, especially in challenging generalization settings such as alignments of unseen ligand binding sites or of catalytic sites - an out-of-distribution setting in which it performed poorly. Another current limitation is its runtime, fast enough for single-query search but not yet for UniProt-scale annotations [Hekkelman et al., 2023]. A clear path to progress is model engineering: scaling-up of the training data (redundant training data, data augmentations and synthetic examples), extensive model architecture and loss hyperparameter search and model distillation should all lead to performance improvements. Extension to other classes of functional sites and protein representations (e.g., surface-based [Gainza et al., 2020, Sverrisson et al., 2021]) could help as well in this regard. Another possible direction is conditional generative models: the model currently outputs a single transformation, but there are multiple valid alignments for flexible ligands or multivalent proteins. Finally, training was only performed on positive pairs; extension to negative pairs, together with end-to-end generation and scoring of alignments, could yield major improvements for database search use cases.

## Acknowledgments

This study was supported by Len Blavatnik and the Blavatnik Family Foundation, the Ministry of Science and Technology (MOST, grant id: 3011006404), the Edmond J. Safra Center for Bioinformatics and the TAD Center for AI and Data Science of Tel Aviv University.

## Author Contributions

H.R., J.T. and H.J.W. conceived the study. H.R. developed the model and performed all the analysis with the help of J.T. H.R. and J.T. wrote the initial manuscript. All authors reviewed and approved the manuscript.

## Data and Code Availability Statement

The data used to train and evaluate LocAlign is publicly available from the BioLIP2 database and the Protein Data Bank. Source code for training, evaluation, and inference of the model is available at https://github.com/hagairavid18/LocAlign

## 4 Methods

### 4.1 Model Architecture

#### 4.1.1 From Protein Structure to Embedded Point Cloud

The first step of the algorithm is to convert the protein structure into a point cloud with per-point 3D coordinates and embedding vector. We work at the *atomic* rather than *residue* resolution because local structural alignments are often only apparent at this resolution. For instance, when aligning two catalytic triads of serine proteases (say, of Subtilisin and Trypsin), the side chains atoms of the catalytic residues would be closely aligned whereas the backbone atoms may not be [Ahern et al., 2025]. There are also cases where there is no one-to-one mapping at the *residue* level. For instance, a large aromatic side chain may be substituted by two smaller hydrophobic side chains. To identify functionally meaningful correspondences, per-atom embeddings must integrate the atom’s intrinsic chemistry and multi-scale spatio-chemical environment with predicted functional annotations, such as an atom’s likelihood of belonging to a ligand-binding site. Importantly, the embeddings must be SO(3)-invariant w.r.t transformation of both the source and target protein to warrant that the correspondences are SO(3)-invariant [Wang and Solomon, 2019]. The embedding extraction transformation should also be identical for both the target and source proteins, to ensure that swapping them yields the same correspondence list.

Based on these considerations, we used the **ScanNet** [Tubiana et al., 2022] **and ESM2** [Lin et al., 2022] models. Briefly, ScanNet is a Geometric Deep Learning model for structure-based functional site prediction. It learns SO(3)-invariant representations of atoms and amino acids based on the spatio-chemical arrangement of the neighbors, and exploits them for residue-level annotation tasks. We used a ScanNet checkpoint pre-trained on a protein binding site prediction task, and extracted the per-atom embeddings, per-residue embeddings and per-residue binding site prediction. We concatenated them together (for a given atom, we stack the per-atom embedding and the per-residue embedding of its parent residue) to obtain a 364-dimensional vector per atom.

ESM2 [Lin et al., 2022] is a family of *sequence*-based Protein Language Models that can be used to extract per-residue embeddings. These embeddings contain a wealth of information about the local structural environment of a residue, and its functional role. Among other applications, Protein language model embeddings were previously shown to be valuable for sequence alignment purposes [Llinares-López et al., 2023, Gonzalez et al., 2024]. Among the ESM2 models, we found that the 18’th layer of `esm2_t30_150M_UR50D` (640-dimensional) offered the best trade-off between performance and runtime. We also tested **ProtTrans** [Elnaggar et al., 2021] **embeddings and found that they consistently underperformed ESM2 in all our experiments. For each atom, we concatenate along the feature dimension the ScanNet and ESM2 features to obtain:**

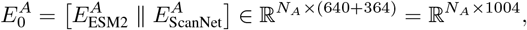

and analogously for 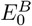 These embeddings are transformed by a multi-layer perceptron (MLP) with shared weights (across atoms and source/target), yielding the final embeddings *E*^*A*^ and *E*^*B*^. In our experiments, we did not attempt to train from scratch or to fine-tune the initial embeddings.

#### 4.1.2 Atom to Atom Edges

In addition to the per-atom coordinates and embeddings, we also calculate atom-atom edge features within the keypoint selection and correspondence solver modules. For each atom *i* of the source or target protein, we first build a reference frame 𝒯_*i*_ ∈ ℝ^4*×*3^, whose center is the atom and whose axes are calculated by Gram-Schmidt orthonormalization of the covalent bonds formed by the atom,following [Tubiana et al., 2022]. Given two atoms *i, j* that belong to the same protein, we define as edge vector the spherical coordinates of atom *j* in frame 𝒯_*i*_ (distance, azimuthal and polar angle). These edge features are invariant upon rigid-body transformation of the protein structure. Importantly, we *do not* include order-dependent features, such as sign(*j* − *i*) to facilitate generalization between order-dependent and order-independent alignments.

#### 4.1.3 Differentiable Keypoints Selection

After calculating per-atom embeddings for each protein, we seek to identify correspondences between them. Naive all-against-all comparison of all atom embedding pairs is (i) compute and memory-intensive, as the number of heavy atoms is large (up to 5K atoms per protein) and (ii) inefficient, since the key correspondences occur within the ligand-binding sites, which are relatively easy to segment. To address this, we introduce a differentiable keypoints selection mechanism that identifies, for each protein, *K* keypoints (with *K* = 1000 in our experiments) - atoms likely to be part of the local alignment. Then, we only compare the embeddings between pairs of keypoints. This is achieved by calculating a per-atom importance score from its embedding using a scalar graph attention module (described below), and then retaining the top-K atoms with the largest scores. Naive application of the top-K selection operation is not differentiable and the standard straight-through estimator of the gradient [Hinton et al., 2012] performed poorly in our experiments. Following the literature on differentiable graph coarsening [Cangea et al., 2018, Gao and Ji, 2019] we achieve differentiability by introducing a gating mechanism. Let:

- *E* ∈ ℝ^*M ×F*^ denote the per-atom embedding matrix,
- *A* ∈ {0, 1} ^*M×M*^ denote the adjacency matrix of the *K*_*neighbors*_-nearest neighbors graph, (*K*_*neighbors*_ = 16 in our experiments)
- 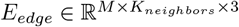 denote the corresponding edge features, calculated as described above.
- *S*^*init*^ ∈ ℝ^*M ×*1^ denote the previous per-atom importance score. In the first iteration, *S*^*init*^ is either uniformly zero or equal to a user-provided motif mask. In subsequent iterations, it is provided by the recycling mechanism.

We subsample the point cloud as follows:

##### Algorithm 1

Top-K Selection Algorithm

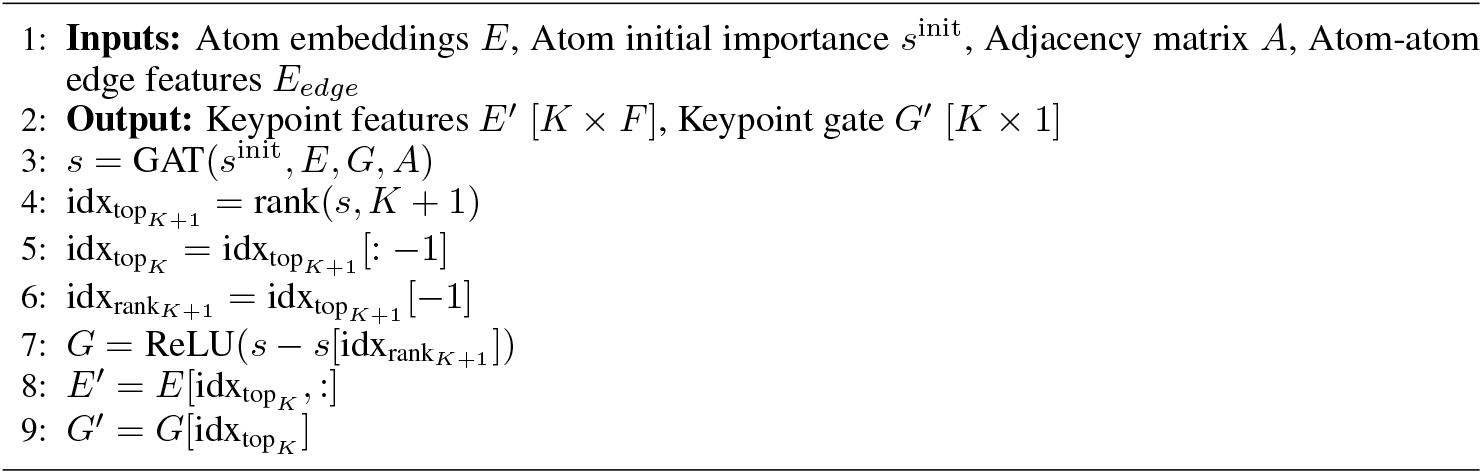

Where GAT is a graph attention layer [Veličković et al., 2017] with scalar output, rank(s,K) is the function that sorts the vector *s* and returns the indices of the top-K entries and g is the learned gate. Unlike prior works [Cangea et al., 2018, Gao and Ji, 2019] that use a sigmoid/tanh activation, we use the ReLU activation with an adaptive bias equal to minus the score of the k+1 atom. By construction, the gate *g* takes zero value for all but the top-K nodes and is differentiable, since the top-K+1 *value* is differentiable. Provided that the subsequent layer is i) Differentiable, ii) Permutation-equivariant and iii) Such that nodes with zero-valued gate do not influence the output 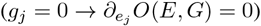, it is easy to see that the array slicing operation (*E*, → *G E*^*′*^, *G*^*′*^) does not change the output, and thus the model is exactly differentiable.

The GAT is implemented as follows. First, we calculate *scalar* keys, queries and values (*q*_*i*_, *k*_*i*_, *v*_*i*_) for each atom using an MLP with three output heads and ReLU output activation function. The initial importance scores are added to the value 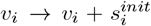. Second, the per-edge vectors are embedded into scalars *e*_*ij*_ as follows. The distance is encoded using Radial Basis Functions (16 learnable kernels, with initial centroids uniformly spaced between 0 and 6 angstroms) and the angles are encoded with sin and cos functions. The 20 features are passed through an MLP with scalar output and no activation function. Finally, the GAT’s output is calculated using the standard attention over neighbors, with the scalar edges used as relative positional encoding:

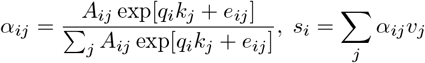

#### 4.1.1 From Embedding to Correspondences

After Keypoint selection, we compute a bidirectionally normalized similarity matrix between the selected atoms, following [Hezroni et al., 2021]. Let:

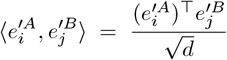

denote the scaled inner product between atom (keypoint) embeddings, where *d* is the embedding dimension. We then define two directional softmax terms, weighted by the learned gate scores

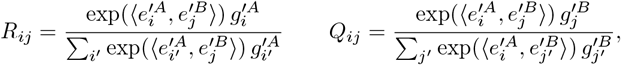

The final soft correspondence matrix is the element-wise product of these two terms:

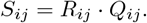

The bidirectional normalization can be intuitively understood as a soft reciprocal best hit (resp. soft best buddy) protocol, following the bioinformatics terminology (resp. computer vision). There is a correspondence between the atoms *A*_*i*_, *B*_*j*_ if the atom that is the most similar in the embedding space to *A*_*i*_, within the protein *B* is *B*_*j*_, and vice-versa. Using the scalar gates 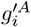 and 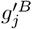 and *g*^*′B*^ ensures that the model is differentiable and biases the probabilities so that more informative atoms are favored in both directions. During training, a dropout with *p* = 0.1 is applied to the soft correspondence matrix, to encourage the model to recover the transformation from a variety of keypoint correspondences.

#### 4.1.5 Correspondence Solver Module (CSM)

The purpose of the Correspondence Solver Module (CSM) is to resolve geometrical inconsistencies in the initial soft correspondence matrix *S* using a Graph Neural Network (GNN). Intuitively, if two correspondences (*i* ↔ *j*) and (*k* ↔ *l*) are true, then the distances between their source points 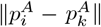 should match the one between their target points 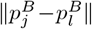. The correct set of correspondences can be thus thought as a maximal clique in a graph where each node is a correspondence and there are edges between geometrically-consistent correspondences. The CSM module uncovers such a clique with a GNN that reinforces correspondences that are mutually consistent and suppresses the others.

Practically, from the (*K* × *K*) soft correspondence matrix *S*, we first extract the top *K*^*′*^ highest-scoring correspondences (*K*^*′*^ = 400 in our experiments) for computational efficiency. To ensure differentiability, we follow Algorithm 1, without using any MLP (*F* = 1, *g* = *E*) and keep only the gates (*g*^*′*^) as a scalar feature.

We next build a graph-based representation where each correspondence (*i* ↔ *j*) forms a node in a fully connected graph. For two correspondences (*i* ↔ *j*) and (*k* ↔ *l*), we take as edge features the concatenation of the atom-atom features 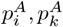 and 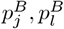, as described above. These features are invariant upon rigid-body transformation of either the source or target protein, and do not depend on the order of the atoms. The distances are encoded using Radial Basis Functions (32 learnable kernels, with initial centroids uniformly spaced between 0 and 30 angstroms). The angles are encoded with sin and cos functions. These 32*2+4*2=72 features are passed through an MLP (without activation function) to produce a scalar edge weight *α*_*mn*_ that quantifies the geometric consistency of the two correspondences. In line with our intuition, we found in our experiments that the *learnt* edge values were large and positive for geometrically-consistent correspondences and negative otherwise.

We obtain an initial graph with scalar node features and edges, to which we next apply *n*_iter_ (*n*_iter_ = 10 in our experiments) rounds of scalar message passing using a GraphConv-style update. Let 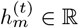 denote the scalar node state for correspondence *m* = (*i* ↔ *j*) after layer *t*. The update rule is

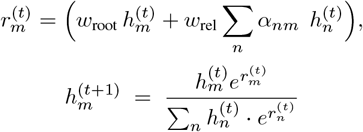

Where *w*_root_, *w*_rel_ ∈ ℝ are learned scalar parameters (shared between layers). The differences from a standard GCN update are that the updates are multiplicative rather than additive, and that the scalars are normalized to one after each iteration. Using multiplicative updates ensures that if a correspondence is initially at zero, it stays at zero, one of our conditions for differentiability after the top-K operation. The final correspondence weights are set to: 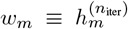. The top *K* correspondences and their associated weights are passed to the transformation estimation module. They provide a compact and geometrically consistent basis for accurate alignment.

#### 4.1.6 From Correspondences to Transformation

We estimate the rigid-body transformation using a *weighted* version of the Kabsch algorithm [Sorkine-Hornung and Rabinovich, 2016], applied only to the top *K*^*′*^ refined correspondences. Each correspondence is assigned a scalar weight from the CSM, normalized such that 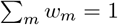. High-confidence matches therefore dominate the alignment, while weak or noisy ones contribute little.

Weighted centroids are computed as:

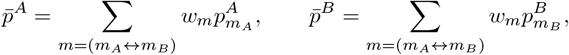

where 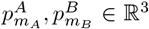 are the coordinates of the corresponding atoms and *w*_*m*_ are the normalized weights.

The weighted covariance matrix *S* ∈ ℝ^3*×*3^ is then:

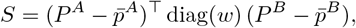

Here, 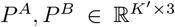 stack the coordinates of the top-*K*^*′*^ correspondences, where each row corresponds to a pair (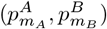) of matched atoms.

We compute the singular value decomposition:

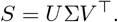

The optimal rotation is:

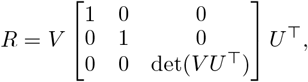

and the optimal translation is:

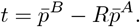

This weighting ensures that the final transformation is determined by a consistent subset of strong correspondences, yielding accurate alignments even in the presence of noisy matches.

#### 4.1.7 Recycling

Classical point cloud registration (RANSAC, ICP) or protein structural alignment algorithms (TM-Align) are iterative: they alternatively estimate the optimal rigid-body transformation from a list of correspondences, and use it to refine the correspondence list. Correspondences between points that are far away after initial transformation are discarded, whereas some initially overlooked correspondences between “passenger” points may become apparent only after a transformation calculated from the keypoints is applied. Prior deep learning-based point cloud registration works also investigated iterative refinement using various heuristics: e.g., DCP [Wang and Solomon, 2019] refines the initial transformation using ICP; GeoTransformer [Qin et al., 2022] runs a RANSAC on the learned correspondences. However, to the best of our knowledge, these refinement protocols are not trained in an end-to-end fashion.

AlphaFold2, [Jumper et al., 2021] introduced a recycling mechanism that enables the model to iteratively refine an initial structure prediction. Briefly, the recycling recipe is as follows: (i) Build a model architecture that takes as extra input an initial guess of the desired output (“template” track). (ii) At the end of each iteration, feed back the output of the model to the initial guess track and use it to generate the next iteration’s output. (iii) Calculate the loss at each iteration; use stop gradients after each iteration to only backpropagate through a single iteration. (iv) Use a fixed number of iterations at training time, and variable at inference time according to the difficulty of the example. Recycling allows to train in a numerically stable fashion an iterative algorithm, with variable number of iterations at inference time.

We follow a similar approach for LocAlign. Intuitively, we first want that atoms that are part of the alignment of the r’th iteration should be used as input motifs for the (r+1)’th iteration. To this end, we calculate the per-atom recycled importance score, defined as the sum of the weights of the correspondences that involve them, times a trainable scaling factor. This score is fed back to the keypoint selection module, as described above.

Additionally, atom pairs that are tightly superimposed after the r’th iteration should get higher initial correspondence score at the (r+1)’th iteration. For (i), a natural representation for the initial guess would be a list of initial correspondences between pairs of atoms 𝒞 = { (*i* ↔ *j*), (*k* ↔ *l*), … }, *stored as a sparse binary array*. Architecture-wise, these initial correspondences would be incorporated as a relative positional encoding for calculating the initial similarity matrix: 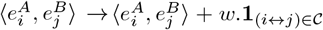. However, it is not obvious how to extract a sparse correspondence array from the superimposed structures without discarding important information. An alternative is to encode the list of initial correspondences using extra per-atom, 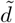 -dimensional positional encoding vectors 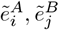, such that 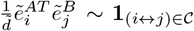 Then, it suffices to concatenate and rescale these vectors with the initial embeddings 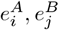, such that the scaled dot product matrix equals the sum of the embedding-based correspondence matrix and of the initial (*dense*) correspondence matrix.

To extract positional encodings from the *superimposed* source and target structures, we use 3D Fourier transforms of their coordinates defined as follows. Let ℱ^*B*^ = (*R*^*B*^, *t*^*B*^) the principal axis frame of the (fixed) target point cloud *P*^*B*^ (or any other SO(3)-equivariant frame). Let 𝒯 ^(*r*)^=(*R*^(*r*)^, *t*^(*r*)^) the rigid-body transformation found after *r* recycling rounds. Let *P*^*A*,(*r*)^ =R^BT^[*R*_(*r*)_*P*_*A*_ + *t*_(*r*)_] − *t*^*B*^ denote the coordinates of the source point cloud, after applying the predicted transformation, in the frame ℱ^*B*^. We detach gradients here (using the .detach()), then calculate the 3D Fourier embeddings as follows:

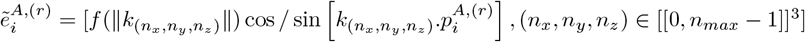

Where 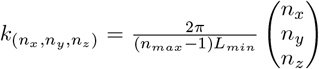 and 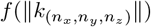 is a trainable scaling function. In our experiments *L*_*min*_ = 5 Å, *n*_*max*_ = 8 such that 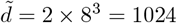. We similarly calculate 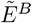, without applying 𝒯 ^(*r*)^.

Using the coordinates in the principal axis frame ℱ^*B*^ warrants that the embeddings are strictly invariant upon rigid-body transformation of either the source or the target protein. The scaling factor 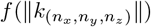 is a trainable function of the norm of the Fourier vector, parameterized using RBFs followed by linear projection (in our experiments, 8 kernels, centered at zeros, initialized with standard deviations 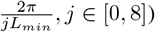). Using scaling factors that are shared between the cos and sin and only depend on the norm of 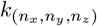 warrants that the dot product between two embeddings 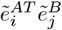 only (approximately) depends on the distance 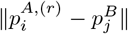. *f* controls the rate at which the dot product decays as function of the distance between the points.

The resulting positional encodings 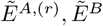 are concatenated with the initial embeddings to calculate the soft correspondence matrix in the next recycling iteration. In our experiments, we used four recycling iterations (i.e., five predictions).

### 4.2 Loss Function and Evaluation Metrics

Our total loss combines a **supervised** ligand alignment term together with an **unsupervised** alignment quality loss, which is has three components. The total loss is defined as:

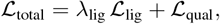

where the quality loss ℒ_qual_ is composed of four parts:

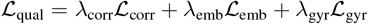

The model checkpoint presented in the article was trained with *λ*_lig_ = 1.0, *λ*_corr_ = 0.2, *λ*_emb_ = 0.1, *λ*_gyr_ = 0.1.

#### Ligand Loss (Supervised)

The supervised component evaluates how well the predicted transformation superimposes the source ligand onto the target ligand. Let 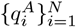 and 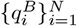 denote the *N* ligand atoms in the source and target structures, respectively. We utilize the atom IDs to establish a one-to-one correspondence between the poses. Let 𝒯 = (*R, t*) denote the predicted transformation. We first compute the raw root-mean-square deviation (RMSD):

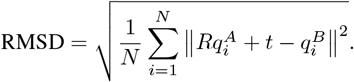

To ensure training stability and numerical robustness, we apply a non-linear transformation to the final loss. The reasoning is that beyond a certain distance *d*_0_, an alignment is considered failed; therefore, the distinction between a “poor” and an “even worse” alignment should not result in an unbounded penalty. The final ligand loss is defined as:

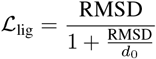

where *d*_0_ = 10.0 Å is a scaling constant. This formulation prevents outliers from dominating the gradient while maintaining a nearly linear behavior for high-quality alignments where the RMSD is small.

#### Correspondence RMSD Loss

To ensure geometric consistency, we define a weighted RMSD over the top *K*^*′*^ correspondences 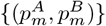 with weights *w*_*m*_ ( ∑*w*_*m*_ = 1). For a predicted transformation 𝒯, we compute the weighted distance:

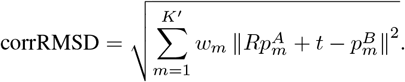

Following the same logic as the *Ligand Loss*, we apply a non-linear normalization for stability: corrRMSD

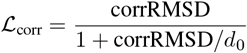

where *d*_0_ = 10.0 Å . This prevents poorly initialized correspondences from dominating the optimization process.

#### Embedding Loss

The purpose of the embedding loss is to encourage correspondences between atoms with high chemical similarity, *i.e*., that can engage in similar physical interactions with ligands (as hydrogen bond donors or acceptors, charges, etc.). A difficulty is that chemical similarity differs from atom type similarity, as e.g., an oxygen atom can be either a hydrogen bond donor (if part of a carboxyl group), hydrogen bond acceptor (if part of an hydroxyl group), negatively charged (depending on the pH), or none of the above (if already engaged in intra-protein interactions). The loss term should thus also encourage the learning of good atomic representations such that proximity in the embedding space indicates similar chemistry.

Let 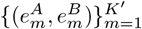 denote the embeddings of the *K*^*′*^ target and source atoms in correspondence, and let 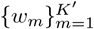 denote the correspondence weights (with 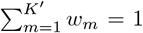). Let *β >* 0 denote a trainable scalar inverse temperature parameter. The loss is given by:

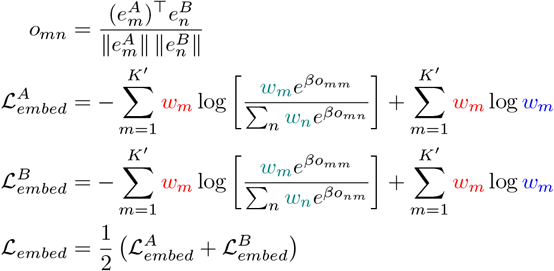

The loss is similar to the standard contrastive loss used in CLIP [Radford et al., 2021], with additional weights injected as described below; for 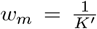, we recover the standard contrastive loss.

- Red weights: The example in the “batch” (= the alignment) are weighted by their respective weights *w*_*m*_, to ensure that correspondences with zero weight do not contribute to the loss (see the condition on keypoint selection).
- Green weights: For the same reason, the examples are also weighted when calculating the softmax distribution.
- Blue weights: We subtract the negative log-likelihood of the null distribution *p*_*m*_ = *w*_*m*_. Without this additional term, the loss can be “hacked” by producing an alignment with a − single correspondence 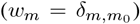. With this additional term, the loss is minimal (= − log *K*^*′*^) if the embeddings are perfectly matched AND if the number of correspondences is maximal 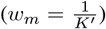. It equals to zero if the embeddings are all identical or *β* = 0, and can be arbitrarily large for mismatched atoms.

Intuitively, an alignment has perfect attribute similarity if ∀*m, n, o*_*mm*_ ≫ *o*_*mn*_ and *o*_*mm*_ ≫ *o*_*nm*_, that is, all aligned atom pairs are reciprocal best hits in the embedding space. Note that we only request that atom pairs are reciprocal best hits within the alignment rather than within the whole protein because the same structural motif could appear twice. The inverse temperature *β* controls the desired margin between corresponding and non-corresponding pairs, and is learned here. Indeed, if it is too large, deviations from reciprocal best hits are heavily penalized, while if it is too small, alignments with multiple correspondences per amino acid are penalized (embeddings of two atoms from the same amino acid are always similar). In our experiments, the learned value of *β* were in the range 40 − 60, in agreement with [Radford et al., 2021].

The dual role of the loss can be understood as follows. When the atomic embeddings are fixed, the loss is minimized by adjusting the weights *w*_*m*_ such that correspondences between atoms with similar (resp. dissimilar) embeddings are reinforced (resp. damped). When the correspondence weights are fixed, the loss is minimized by adjusting the embeddings such that atom pairs in correspondence should have similar embeddings. This “chicken-and-egg” process does not collapse to a trivial solution because the correspondences are constrained by the rigid 3D geometry: two atoms with low embedding similarity can have a high correspondence weight because they are well-superimposed in 3D.

#### Radius of gyration loss

At *fixed alignment size*, alignments that are more compact are more likely to be biologically significant than alignments that are scattered. The radius of gyration loss is designed to encourage compact alignments, without biasing the alignment size. It is defined as:

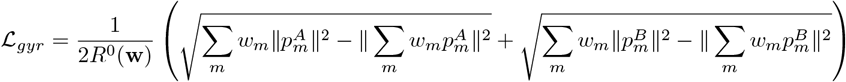

Where *R*^0^(**w**) is a reference radius of gyration that scales with the effective number of correspondences 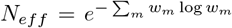. In our experiments, we used 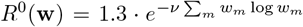 and *ν* = 0.4 following standard scaling laws for protein structures [Hong and Lei, 2009].

#### Atom matching score metric

To evaluate the quality of the correspondences, we explicitly test whether matched atoms belong to the same chemical type. Since a meaningful alignment should preserve atom identities, we define a consistency score *S*_*type*_. In protein structures, the vast majority of heavy atoms are Carbon; consequently, a simple unweighted matching score would be expected to be very high regardless of alignment accuracy. To account for this non-uniform distribution, we introduce importance weights Ω(*T*) that are approximately inversely proportional to the frequency of each atom type (C: 1, O: 3, N: 4, S: 100).

Let 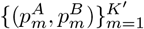 denote the *K*^*′*^ correspondences with weights 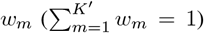, and let 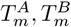 be the atom types for the source and target atoms. We define the importance of a match as the average weight of the two types: 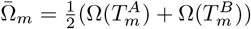. The final score is calculated as:

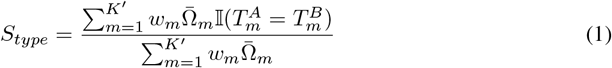

where 𝕀(·) is the indicator function. This formulation ensures that correctly matching chemically distinct atoms like Sulfur contributes more to the metric than Carbon-Carbon matches.

To provide context for this metric, we calculate a random expectation baseline based on the background distribution of atom types within the full protein structures. Let *P*_*A*_(*i*) and *P*_*B*_(*i*) be the probability of occurrence for atom type *i* in the source and target proteins, respectively. The expected numerator is defined as ∑_*i*_ *P*_*A*_(*i*)*P*_*B*_(*i*)Ω(*i*), while the expected denominator is the mean of the expected weights: 0.5 ×( ∑_*i*_ *P*_*A*_(*i*)Ω(*i*)+ ∑_*i*_ *P*_*B*_(*i*)Ω(*i*)). Under this weighted formulation, the random expectation value is approximately 0.45. Consequently, we set the threshold for chemical consistency at *S*_*type*_ *>* 0.5, serving as one of the criteria used to define a successful alignment.

### 4.3 Alignment Quality and Model Confidence Scores

After computing the alignment, we report the following five alignment quality metrics:

- The correspondence RMSD ℒ_*corr*_.
- The perplexity *N*_*eff*_ = exp[ −∑_*m*_ *w*_*m*_ log *w*_*m*_], which can be interpreted as the effective number of correspondences.
- The attribute embedding similarity score: 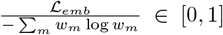. It equals to one if the corresponding atoms are perfectly matched in the embedding space, irrespective of the perplexity.
- The (unnormalized) radius of gyration: 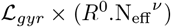.
- The atom matching score *S*_*type*_ as described above.

We define the confidence score as the predicted LRMSD (pLRMSD), which is calculated from the first four above metrics by a monotonic Generalized Additive Model (GAM). A monotonic GAM is a predictive model of the form *y* = ∑_*j*_ *f*_*j*_(*x*_*j*_), where *x*_*j*_ are the features, *y* is the target, and *f*_*j*_ are trainable, monotonic non-linearities. The GAM was trained on the validation set, using the ligand RMSD as target, clipped between 0 and 10 and the mean absolute error loss. The GAM was implemented using scikit-learn’s HistGradientBoostingRegressor with no interactions and monotonicity constraint. The gradient boosting’s hyperparameters (*L*_2_ regularization, learning rate, number of boosting iterations and maximum number of leaf nodes) were selected by cross-validation, using a per-ligand group split and the Spearman correlation coefficient metric.

### 4.4 Data

#### Database creation

We built our database from the BioLip 2025-07-30 version, which contains 989,724 entries. Each entry specifies a PDB chain and the ligand chain to which it binds. To ensure that the ground truth transformation is unique, we first filtered out protein chains that bind multiple copies of the same ligand. Following [Abramson et al., 2024], we also excluded ligands commonly used as crystallization aids and promiscuous binders (Supplementary Table 10 of [Abramson et al., 2024]). After filtering, we were left with 1940 unique ligands bound to 22 467 unique protein chains.

#### CATH Similarity Degree

The complexity of protein alignment is inherently tied to the evolutionary relationship between the structures. To quantify task difficulty, we utilize the CATH hierarchical classification system [Waman et al., 2024], which organizes protein domains into four primary levels: Class (C), Architecture (A), Topology (T), and Homology (H). For a given pair of protein chains, we define the **CATH similarity degree** as the highest level of hierarchical agreement between any two constituent domains.

Specifically, the similarity degree *d* ∈ { 0, 1, 2, 3, 4 } represents the depth of the shared CATH prefix. For example, a similarity degree of 3 indicates that the most similar domain pair shares a common Topology (e.g., both domains belong to 3.40.50), but does not share a common Homology class. To ensure the model learns to align structures with significant geometric and evolutionary variation, we include pairs with a maximum similarity degree of 4 (Homology), thereby excluding near-identical functional families that would trivialise the alignment task.

#### Pair preparation

Theoretically, for each ligand that binds *n* chains, we can form up to 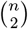 chain– chain pairs; however, doing so would create a highly imbalanced dataset, with an over-representation of (i) some ligands that binds thousands of chains and (ii) easy pairs to align, that have high similarity between examples. To maintain balance, we discard all pairs with CATH similarity equal or above to 5 (i.e., if two of their domains have at least 35% sequence identity), and keep at most 10,000 pairs per ligand, prioritizing examples with lowest CATH similarity degree. Overall, our database contained approximately 140,000 protein–protein pairs, divided into five categories corresponding to the CATH similarity levels 0–4 (see Supplementary Fig. S1).

### 4.5 Database search

#### Template Database

We first created a non-redundant version of the BioLIP2 database as follows. We discarded all BioLIP2 entries that satisfied one of the above:

- The ligand is DNA, RNA, peptide, unknown, or belonged to list of excluded ligands from AlphaFold3.
- The resolution is larger or equal to 4 Å or there is no resolution information available.
- The chain ID is not a single character (e.g., “ab”,…).
- The sequence length is larger than 1024 amino acid.

Next, we clustered the sequences using MMSeqs2’s easy-cluster utility, with a 90% sequence identity threshold and a 90% coverage threshold (cov-mode 0). We retained one representative per cluster, selected as the one with best resolution. We ended up with ∼ 33 000 templates.

#### Template motif

We tried both protein-protein database and protein-motif database search. For each template, we defined the ligand-binding residues as the one for which at least one heavy atom is within 4Å of one heavy atom of the ligand. For the protein-motif comparison experiment, the motif atoms were defined as all atoms that belong to a ligand-binding residue.

#### Hit scoring and ranking

For the protein - protein database search, the pLRMSD score described above is used as the main quality metric (lower is better). For the protein - motif database search, we observed that the pLRMSD score was biased towards small ligands, as these tend to be easier to align. To address this, we calculated for each template an expected LRMSD (eLRMSD) score from the number of atoms in the bound ligand. eLRMSD is obtained by fitting a monotonic GAM on the validation set, using a similar methodology as the pLRMSD. The mean absolute error loss was replaced by a quantile loss with q=0.2, to get the expected value of successful alignments. eLRMSD values ranged from ∼ 1.2 Å for single-atom ligands to 3 − 4 Å for large ligands. The final ranking metric is defined as the ratio pLRMSD/eLRMSD (lower is better).

#### Hardware requirements and runtime

The runtime is reported for a slurm job with 32 CPU threads, 128GB of RAM and one L40S GPU with 48GB of high-bandwidth memory. The database search consists of two stages: a preprocessing stage (run once per database), and the search stage (run once per query). In the preprocessing stage, protein structures are downloaded and parsed, and ScanNet, ESM2 embeddings are pre-computed. It was completed in less than three hours, and generated ∼ 100 GB of cached files in total. The search stage consists in running the ∼ 23, 000 LocAlign inference steps, calculating pLRMSDs, sorting the results, and generating the output ChimeraX scripts for the top hits. It completed in 15-30 minutes, depending on the query size.

### 4.6 Baseline Implementation Details

We compared LocAlign against six established baselines.

- **TM-align** was executed using the `tmtools (v0.2.0)` package, with the returned superposition RMSD used as the correspondence metric.
- **DALI** alignments were generated using the `DaliLite.v5` executable; we automated the pipeline by first converting PDBs to `.dat` format via `import.pl` and then running `dali.pl` with the `--outfmt transrot` flag to extract the optimal transformation matrices and Z-scores.
- **APoc** [Gao and Skolnick, 2013a] was implemented by utilizing `fpocket (v4.2.2)` [Le Guilloux et al., 2009] for the initial extraction of candidate binding sites. To ensure comprehensive coverage, we performed an exhaustive pairwise comparison between the top 10 pockets of the source and target structures, selecting the alignment that yielded the maximum PS-score. In cases where no valid alignment between any pocket pairs existed, the global structural alignment was used.
- **US-align** (v20241108) was evaluated across three distinct mapping modes: the default sequential alignment, fully non-sequential (`fNS`, `-mm 5`), and semi-non-sequential (`sNS`, `-mm 6`). The semi-non-sequential configuration gave the best performance in all datasets and is reported in the main table. We constrained the algorithm to protein-specific atom selection (`-mol prot`) and exported the optimal rotation matrices using the `-m` flag to facilitate coordinate transformation.
- **SoftAlign**. We tested both pretrained checkpoints without and with Smith-Waterman and found that the checkpoint with Smith-Waterman gave the best results (`models/CONT_SW_05_T_3_1`). SoftAlign returns a similarity (soft correspondence) matrix but no 3D transformation. We calculate the latter from the similarity matrix by applying the weighted Kabsch algorithm described above. .
- **PLASMA**. The checkpoint was trained following the instructions from the Github repository: using a Siamese network (hidden dim: 512), a 1,500-pair training subset, Sinkhorn temperature *τ* = 0.1, kernel size *K* = 10, and a matching threshold *ρ* = 0.5.

### 4.7 Additional Implementation Details

For each protein structure, we limit the input to 5, 000 atoms and *during training*, we further randomly subsample 2, 000 atoms, applying padding where necessary. Input features are processed by an initial embedding module of two sequential *FeatureBlocks*. Each block follows a residual architecture using a two-layer MLP (hidden dimension 256) with ReLU activations and dropout (*p* = 0.1), followed by LayerNorm. An identical two-block architecture is used in the *KeypointSelection* module to generate 3-dimensional features (key, query, and value) for the Graph Attention Network (GAT).

The *EdgeWeightLearner*—a residual 2-layer MLP with BatchNorm and ReLU—is utilized twice to compute edge scalars: once for the GAT in keypoint selection, and once for a GNN in the Correspondence Scoring Module (CSM). Based on these scores, the top 400 correspondences are selected for the final alignment.

The total loss is a weighted sum of four components: embedding consistency (*λ* = 0.1), ligand RMSD (*λ* = 1.0), radius preservation (*λ* = 0.1), and correspondence RMSD. To stabilize the geometric refinement, the correspondence loss weight is modulated by a quarter-sine scheduler, ramping from 0.01 up to 0.1 over the course of training.

The model contains approximately 1.6 million trainable parameters. Training is performed using the Adam optimizer with a OneCycle learning rate scheduler, which modulates the learning rate up to a peak of 0.005 with a divide factor of 25. Using a batch size of 8 on a single NVIDIA H100 GPU, training converges within 100 minutes. Inference for 2, 320 samples requires approximately 42 seconds ( ∼ 0.018s per sample), assuming pre-extracted ScanNet and ESM embeddings. All experiments were implemented in Python 3.10.19 using PyTorch 2.9.0 and PyTorch Lightning 2.5.6.

## Supplementary Information

### Data Distribution

**Figure S1.**
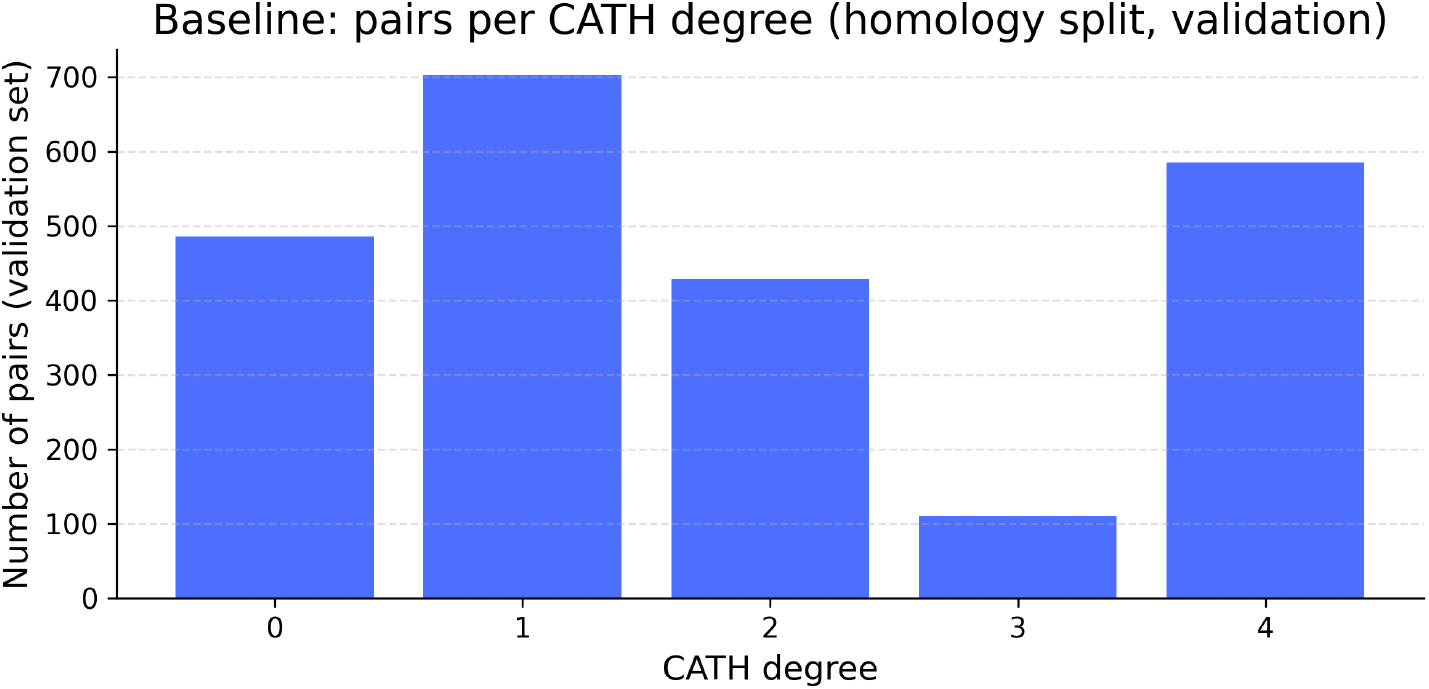
Number of examples per CATH degree. Lower CATH degrees correspond to greater evolutionary divergence and thus more challenging alignment tasks.

#### Ablation study of loss components

We performed an ablation study to investigate the importance of the two main loss terms in the *LocAlign* objective function: the quality loss and the ligand RMSD loss. The quality loss is responsible for the geometric tightness of the alignment, ensuring low correspondence RMSD, consistent atom-type correspondences via embedding loss, and spatial concentration through the radius loss. The ligand RMSD loss ensures that the resulting alignment is functionally centered on the ligands themselves.

**Figure S2.**
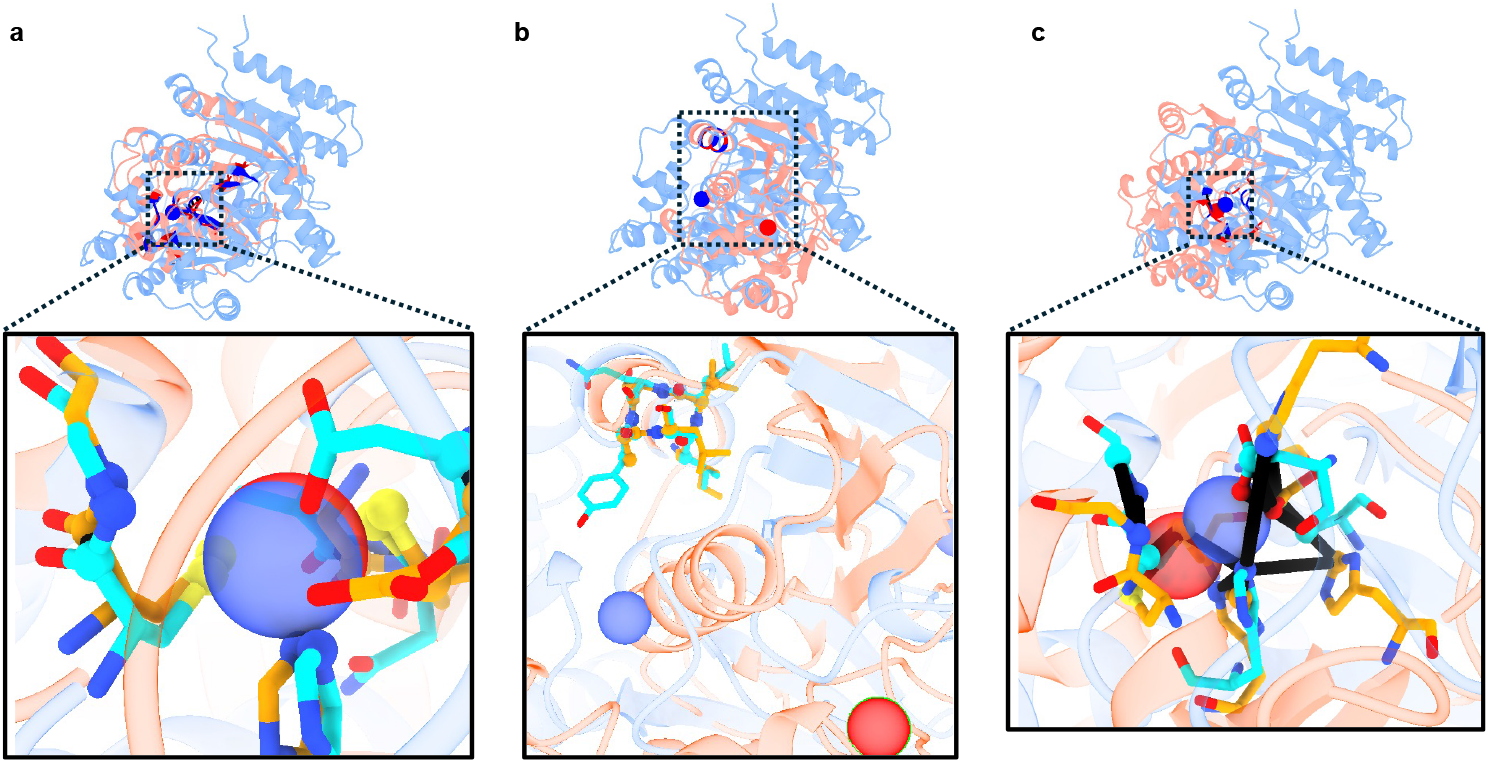
Ablation study of the LocAlign loss components (Example: 1ddz:A ↔ 7bez:A, Zn^2+^). (A) Full model utilizing both quality and ligand loss, resulting in high-fidelity correspondences and aligned ligands. (B) Model without ligand loss; the alignment shifts to a shared alpha-helical motif, leaving the ligands unaligned. (C) Model without quality loss; the ligands are aligned, but correspondences exhibit high RMSD and poor chemical (atom-type) consistency.

#### Inference visualizations

The following examples provide additional evidence of *LocAlign*’s ability to identify conserved structural motifs across proteins with disparate global topologies.

#### Linear motif alignments

We first examine further instances of linear motif conservation (Supplementary Fig. S3):

- **Panel a: 1ci3:M** ↔ **1co6:A (Ligand: HEC)**. This alignment focuses on the Heme C (HEC) binding site. Despite the proteins exhibiting entirely different global folds, the HEC ligand is perfectly superimposed. The model identifies tight, high-fidelity atomic correspondences within the local binding environment, demonstrating its robustness in extracting local functional sites from non-homologous scaffolds.
- **Panel b: 7ph3:A** ↔ **7x4n:E (Ligand: ANP)**. This case highlights the model’s performance with Phosphoaminophosphonic Acid-Adenylate Ester (ANP), a flexible ligand. While the ligand’s global conformation may vary, *LocAlign* identifies precise correspondences concentrated around the phosphate tail of the ligand and their respective protein contact points.

**Figure S3.**
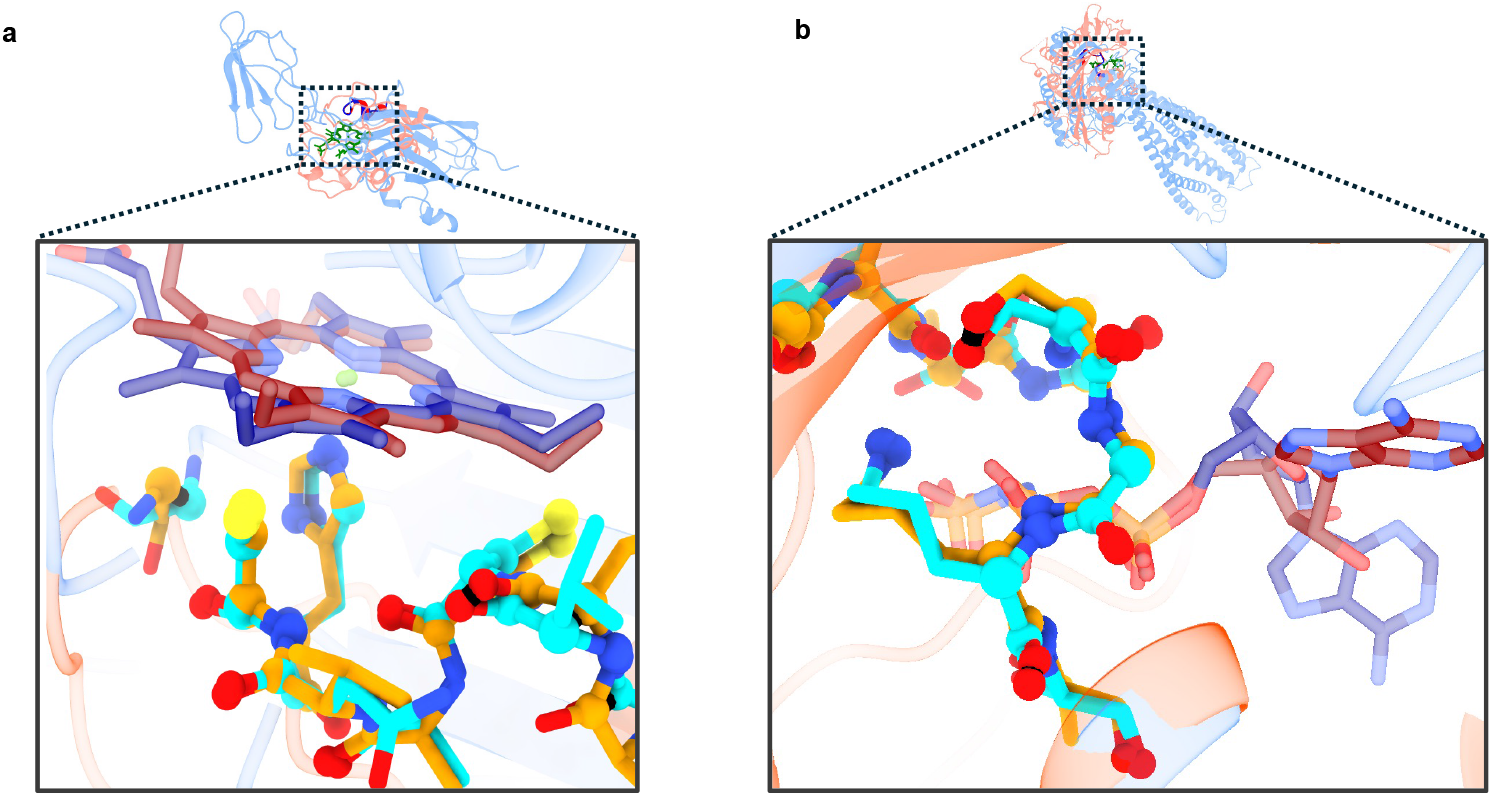
Additional visualizations of linear motif and ligand binding site alignments. (A) Local alignment of Heme C binding sites across divergent folds. (B) Atomic correspondences within the binding pocket of the flexible ANP ligand.

#### Polymorphic and Partially Conserved Chelation Sites

Supplementary Fig. S4 illustrates how the model handles chemical substitutions and complex geometric rearrangements in ion-binding sites:

- **Panel a: 4oj8:B** ↔ **5zur:B (Ligand: FE2)**.
- **Panel b: 1zaa:A** ↔ **1ptq:A (Ligand: ZN)**.
- **Panel c: 5nns:A** ↔ **6ryx:A (Ligand: CU)**.

**Figure S4.**
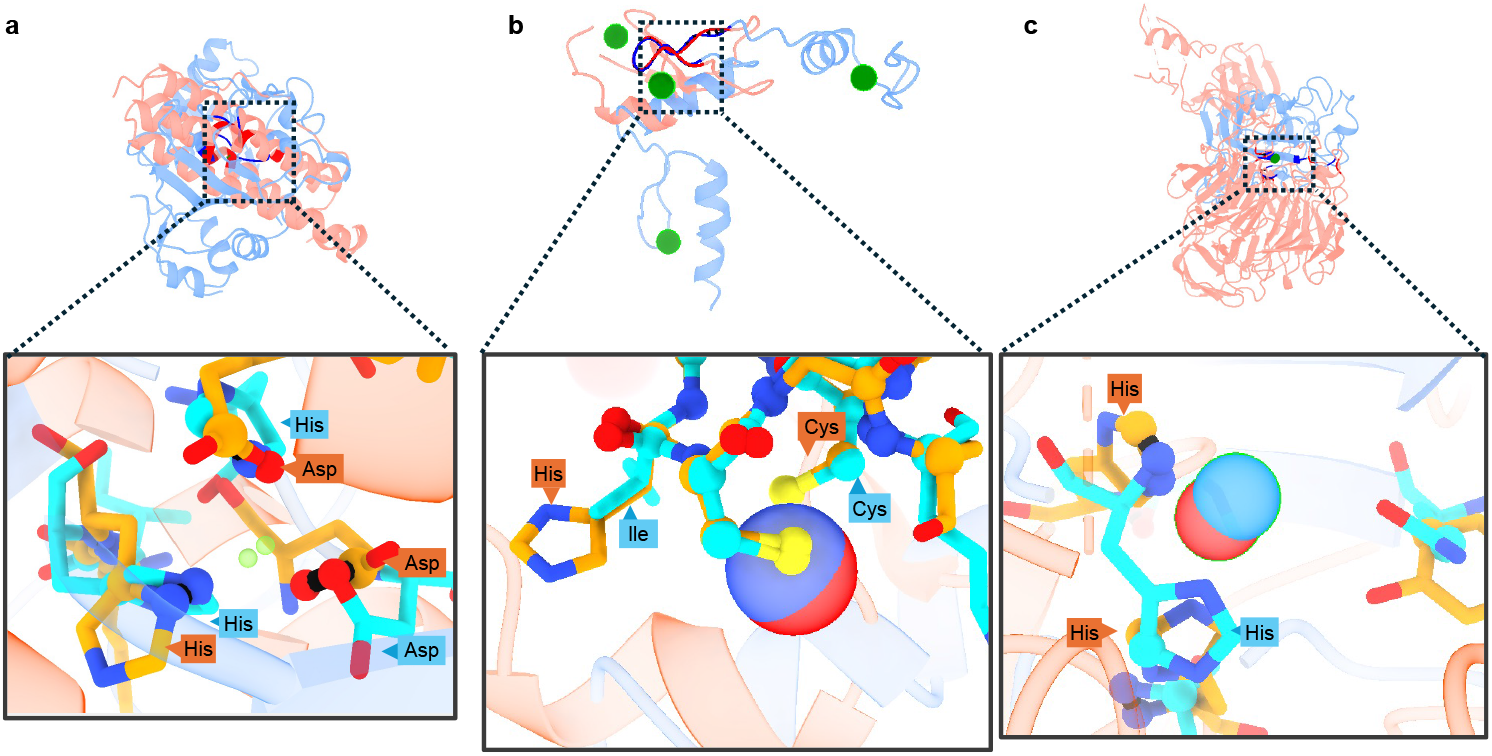
Partially conserved and polymorphic chelation sites. (A) Polymorphic iron-binding site. (B) Zinc-binding site alignment excluding a non-conserved Cys-His substitution. (C) Copper-binding site exhibiting a “two-to-one” substitution where a single histidine replaces the coordination geometry of two distinct histidines.

#### Ground-truth discrepancies and high CATH degree failures

Supplementary Fig. S5 illustrates cases where *LocAlign* yielded high-confidence alignments that were technically classified as failures based on the ligand RMSD threshold (*>* 4 Å). These instances often highlight problematic ground-truth annotations or incomplete structural contexts rather than algorithmic deficiencies:

- **Panel a: 4ymu:J** ↔ **5o6m:D (Ligand: ATP)**. The model identified a high-confidence local alignment anchored by a linear motif containing an alpha helix. However, because the dataset defines a more global structural solution as the “optimal” reference, the ligands are not superimposed in this specific local frame.
- **Panel n: 2dcl:A** ↔ **4ini:A (Ligand: AMP)**. This case demonstrates the impact of **incomplete receptor context**. The biological assembly of the AMP receptor is multimeric; however, the input provided to the model consisted of isolated chains. Consequently, the resulting binding pocket is incomplete, lacking the full suite of chemical signals and steric constraints required to retrieve the optimal alignment that would superimpose the ligands.
- **Panel c: 1hm8:A** ↔ **4isx:A (Ligand: ACO)**. These protein chains appear to possess two distinct potential binding sites. In the source structure (1hm8), the Acetyl-Coenzyme A (ACO) ligand occupies one site, while in the target (4isx), it is bound at the secondary site. Although the protein-level alignment is geometrically sound and shows significant structural overlap, the divergent binding locations result in an RMSD *>* 4 Å, leading to a technical failure classification.

**Figure S5.**
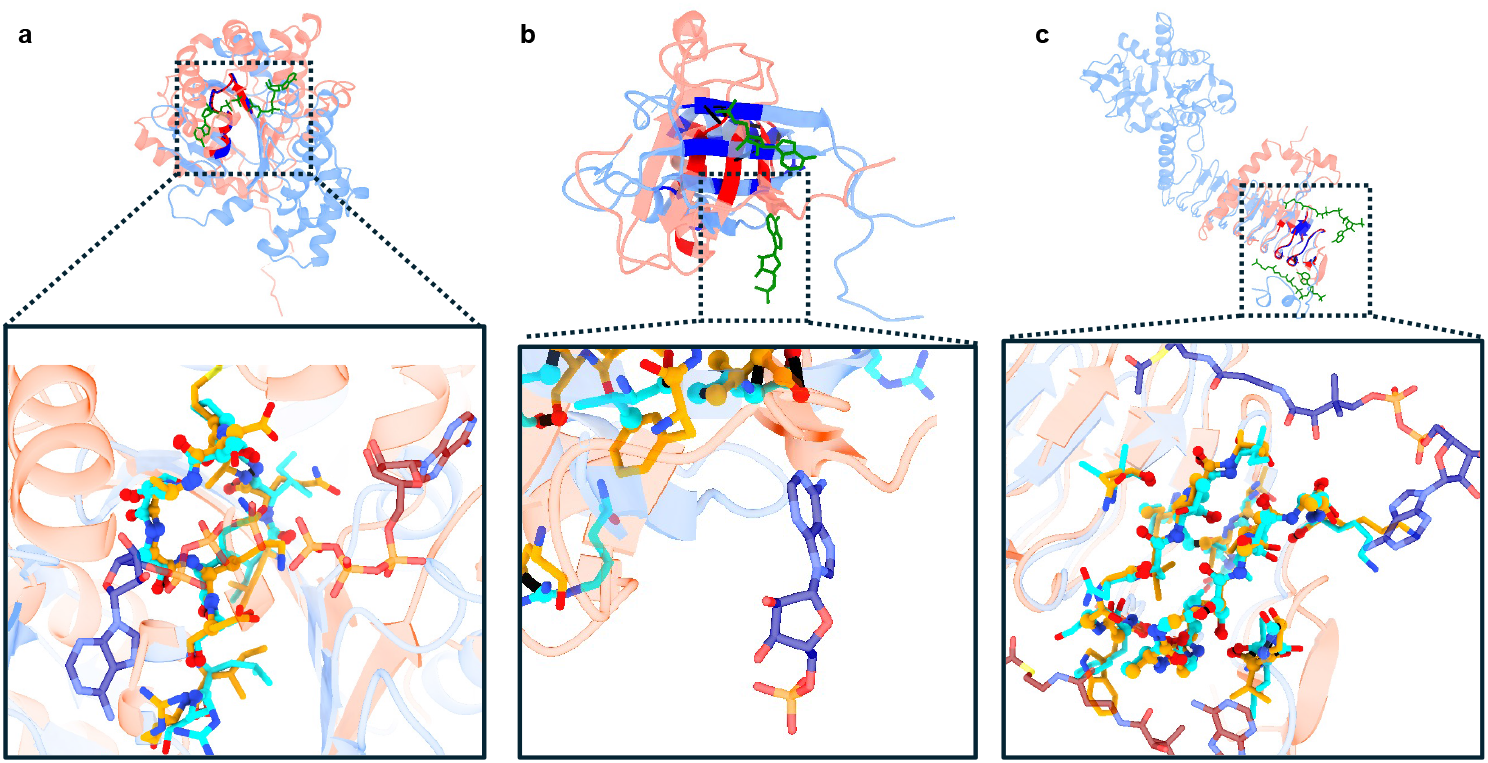
Analysis of high CATH degree failures and problematic ground-truth examples. (a) Preference for an alternative local motif over the annotated global solution. (b) Incomplete binding pocket representation due to missing subunits in a multi-chain receptor. (c) Discrepant ligand binding sites (occupying different “phases”) across the compared structures.

#### Failures in low CATH similarity samples

Supplementary Fig. S6 presents instances where the model failed to retrieve the ground-truth alignment for pairs with low CATH similarity. These cases represent a significantly more challenging task, as the structural signal for alignment is substantially weaker. The model relies on identifying robust geometric and chemical patterns to construct correspondences; however, in these examples, the binding pockets do not exhibit enough similarity to reconcile the divergent scaffolds with the annotated ground-truth.

**Figure S6.**
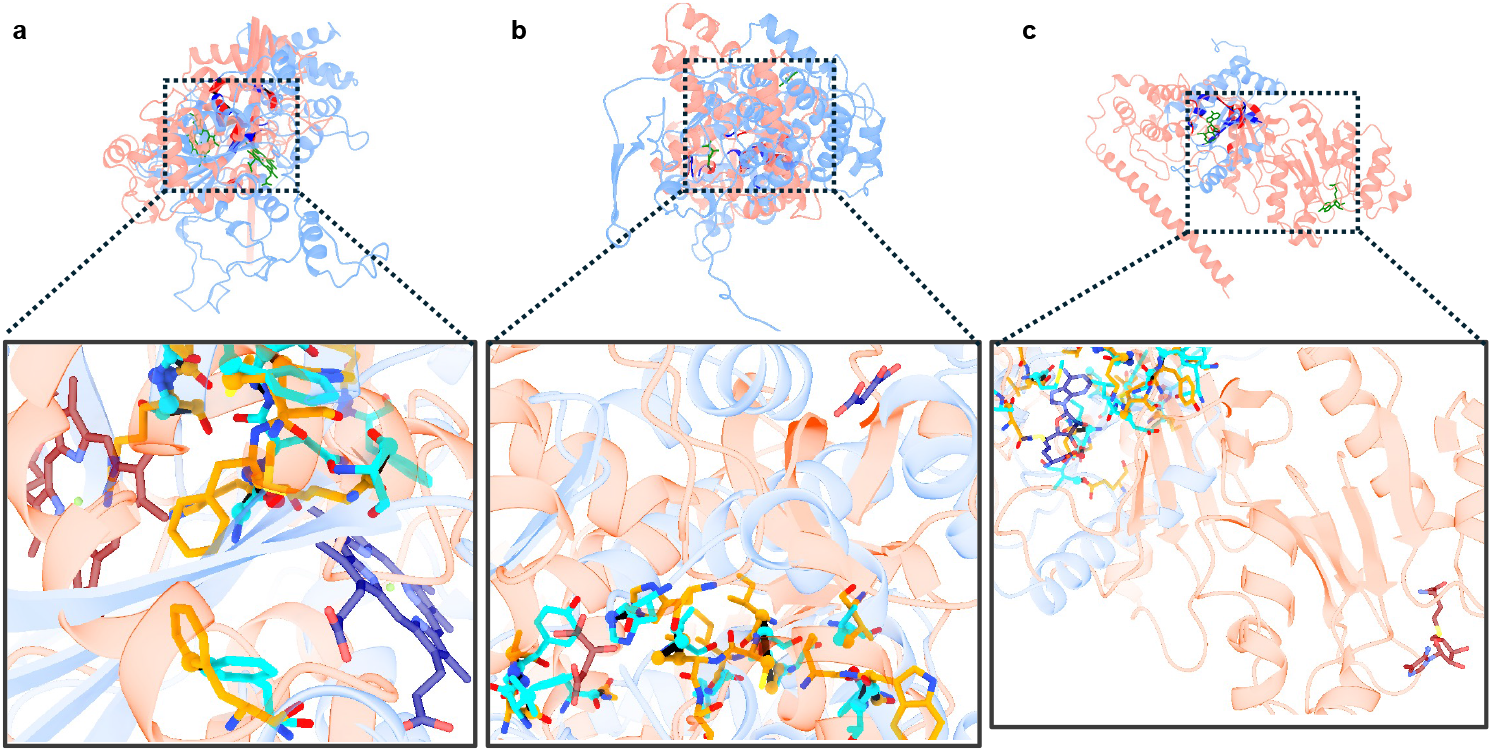
Characterization of failures in samples with low CATH similarity. (a) 1si8:A ↔ 4cuo:A (HEM), CATH degree 0. (b) 3msu:B ↔ 4ros:A (OAA), CATH degree 0. (c) 4fak:A ↔ 4uw0:A (SAM), CATH degree 2.

#### Performance vs. CATH similarity

As shown in Fig. S7, performance is notably higher at CATH similarity degree 4 (same homologous superfamily). However, across degrees 0–3, we do not observe a monotonic trend. This suggests that once global folds differ (degrees ≤ 3), local alignment performance is largely decoupled from global structural similarity.

**Figure S7.**
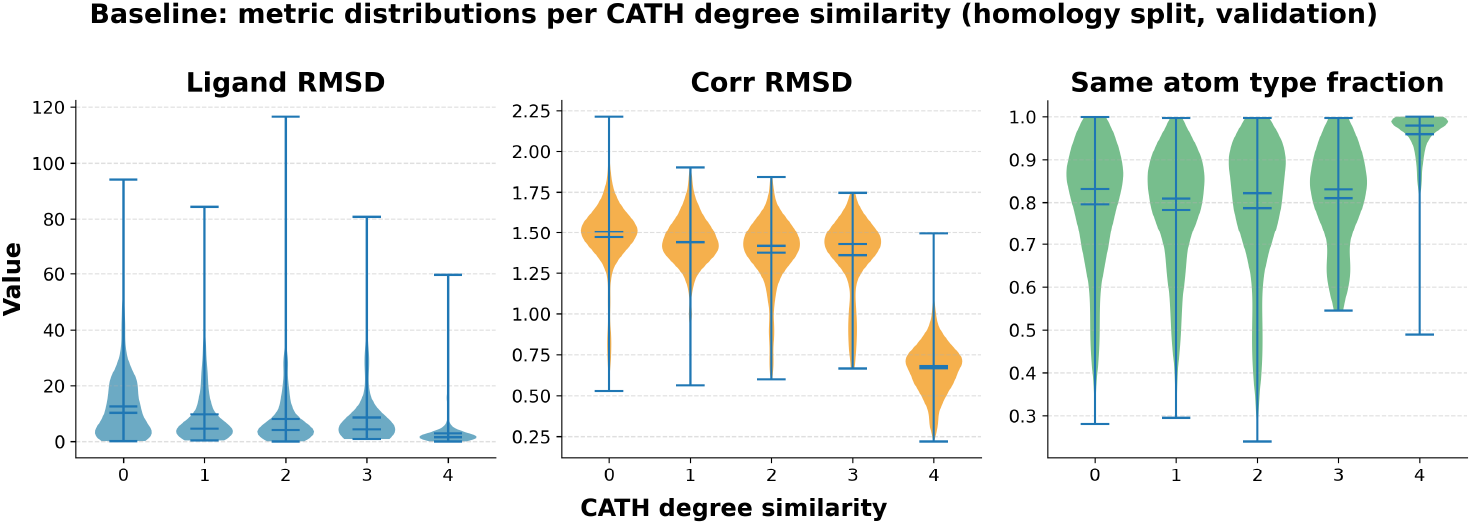
Distribution of alignment metrics (Ligand RMSD, Corr RMSD, Atom type fraction) across CATH similarity degrees. Lower RMSD values and higher atom type fractions indicate better alignment quality.

#### Model Confidence (pLRMSD)

**Figure S8.**
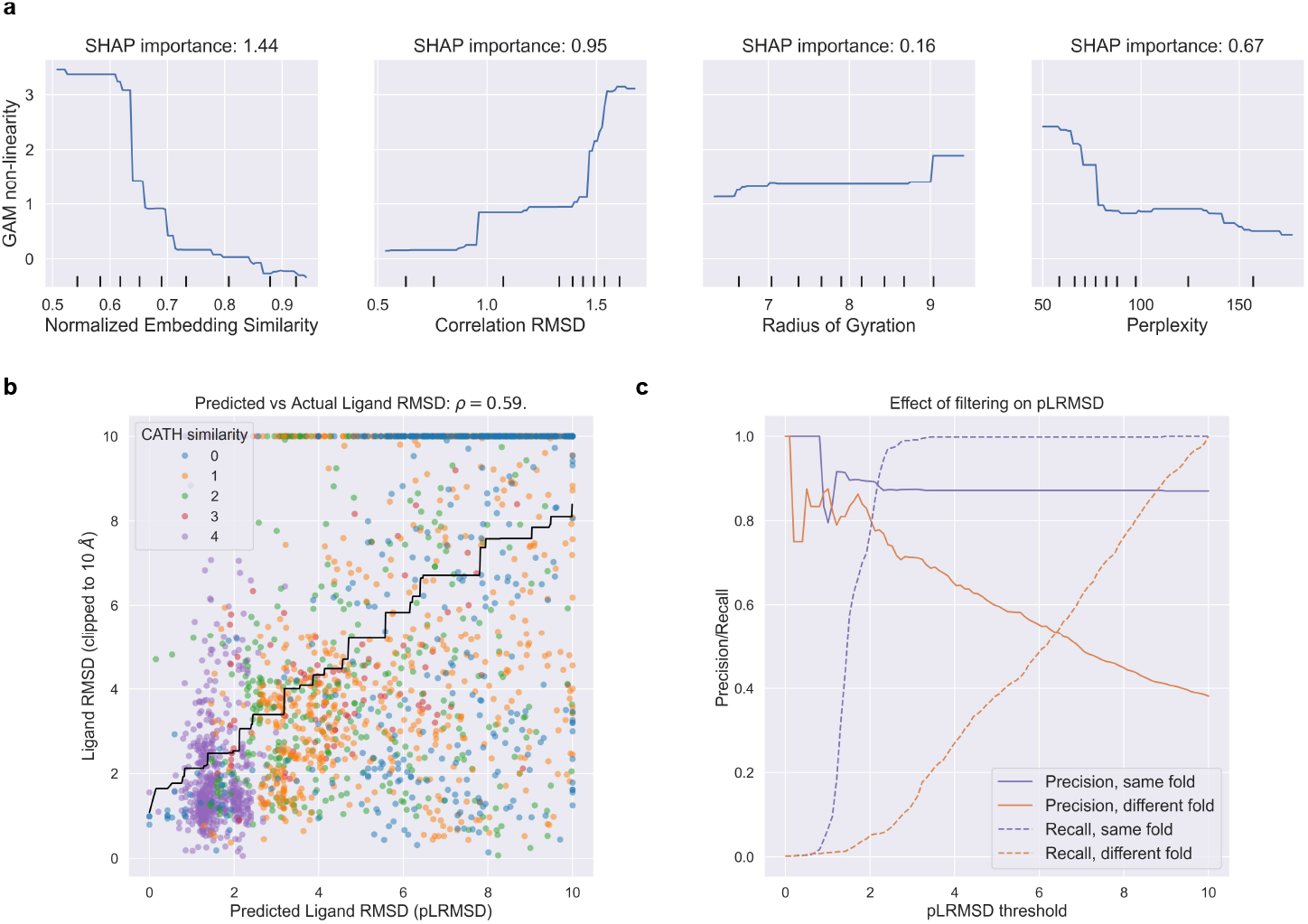
Model Confidence Prediction Head). The confidence is defined as the predicted Ligand Root Mean Square Deviation (pLRMSD), and is calculated with a monotonic GAM: pLRMSD = ∑_*j*_ *f*_*j*_(*q*_*j*_), where *q*_*j*_ are the alignment quality scores and *f*_*j*_ are trainable monotonic non-linearities. (a) The learned non-linearities *f*_*j*_ for each quality score (b) Scatter plot comparing predicted versus actual Ligand RMSD on the validation set. Predictions are made in a cross-validation setting, using a group split on ligands and LRMSD values are clipped at 10Å . The black solid line depicts the isotonic regression fit. (c) Filtering alignments by pLRMSD. For each pLRMSD threshold *θ*, we report the precision (success rate over alignments with pLRMSD ≤ *θ*, full lines) and the recall (fraction of alignments with pLRMSD ≤ *θ*, dotted lines), for pairs with similar (purple) and different (orange) fold.

#### Database Search

**Figure S9.**
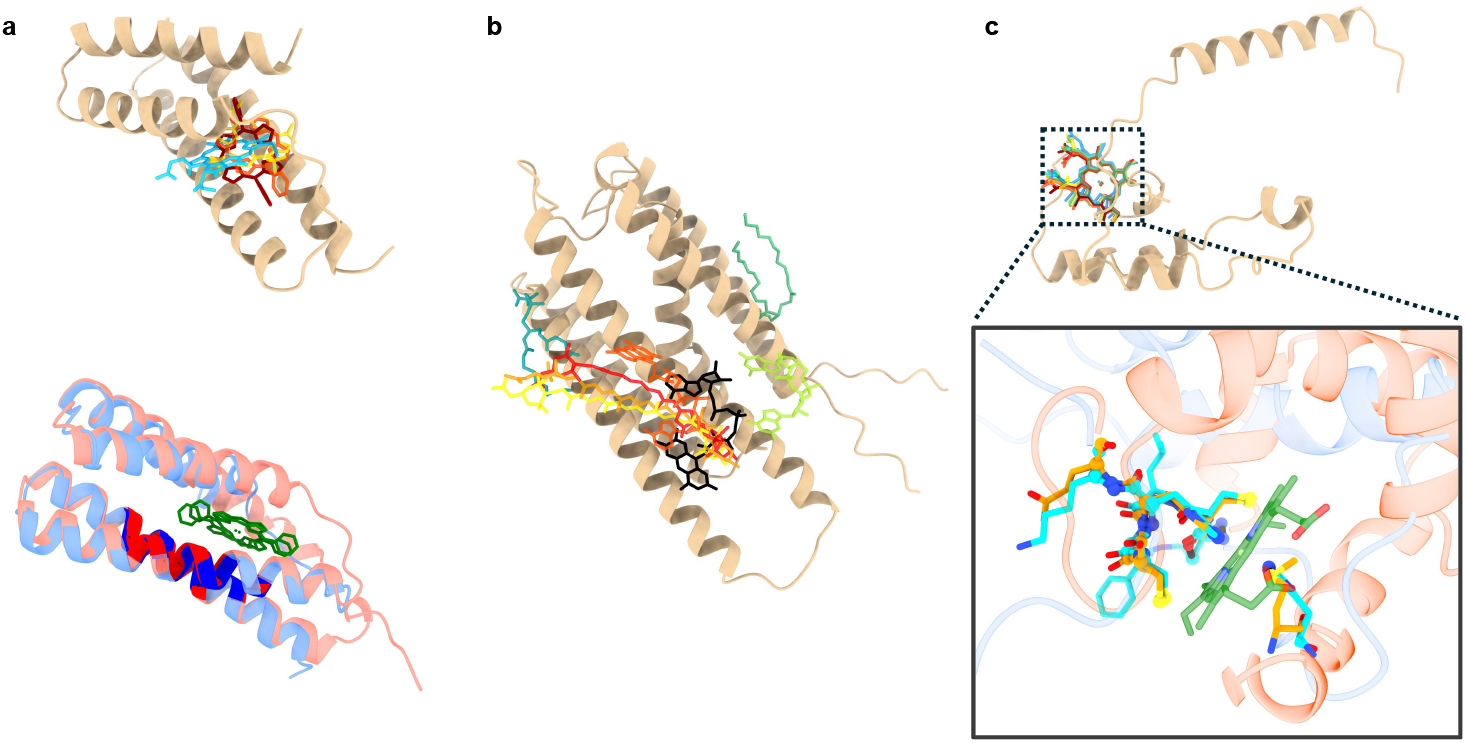
Performance of LocAlign in database search queries and low-confidence hits. **(a) Retrieval of structural homologs:** Querying the de novo designed porphyrin-binder 9cte:A retrieves its architectural progenitor (template 7jrq:A, pLRMSD = 1.83Å). LocAlign identifies the conserved structural motif and relative positioning between the template (WUP) and query (SMU) ligands. (b) **Low-confidence search results:** Example of a query returning poor structural correspondences. All top hits exhibit a pLRMSD *>* 4Å, indicating a lack of significant structural similarity to known BioLIP2 binding sites. (c) **Cytochrome C-binding motif identification:**Alignment of a predicted heme binder against a Heme C (HEC) site (template 3hq9:A, pLRMSD = 0.79Å).

**Table S1.**
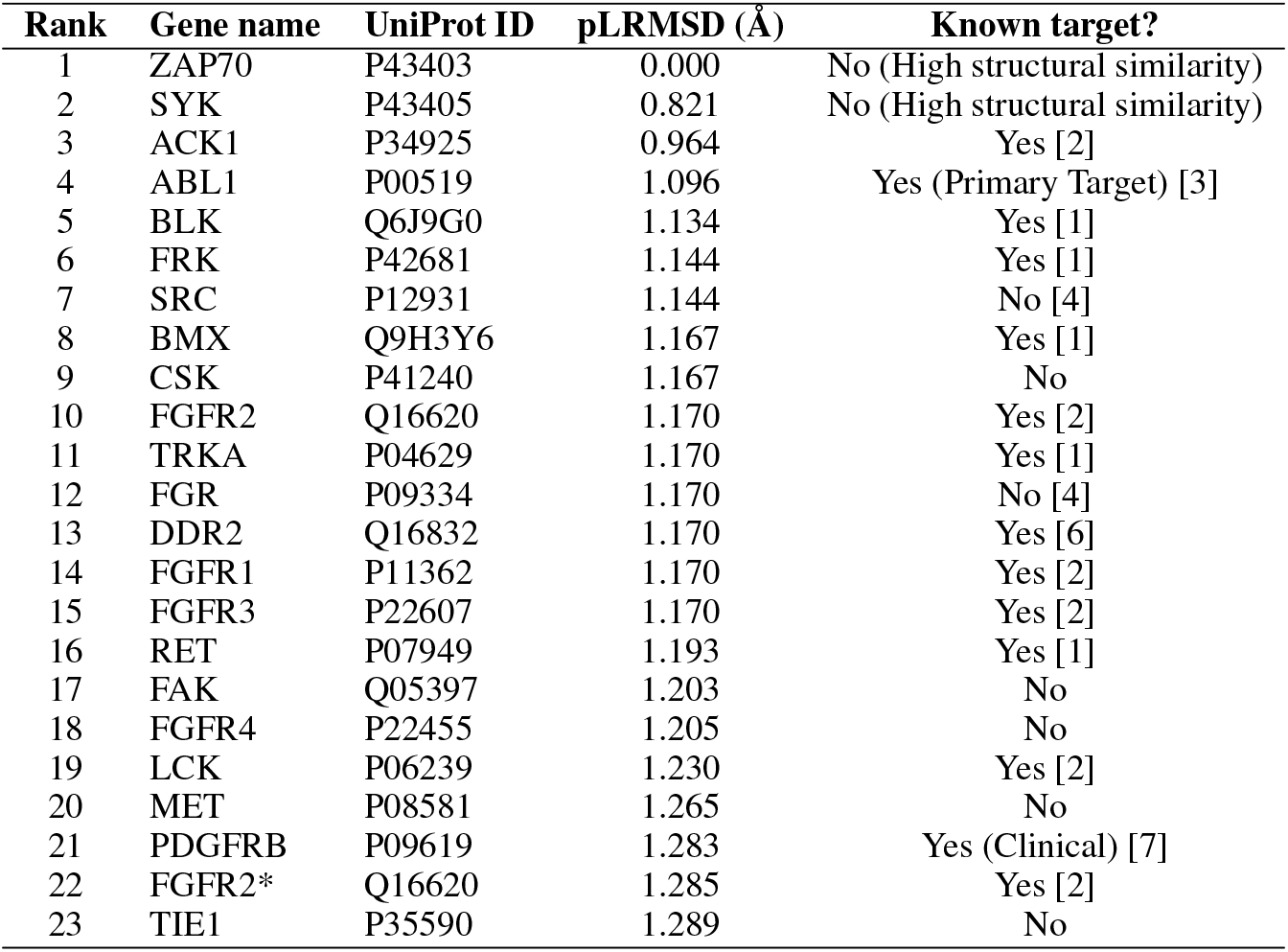
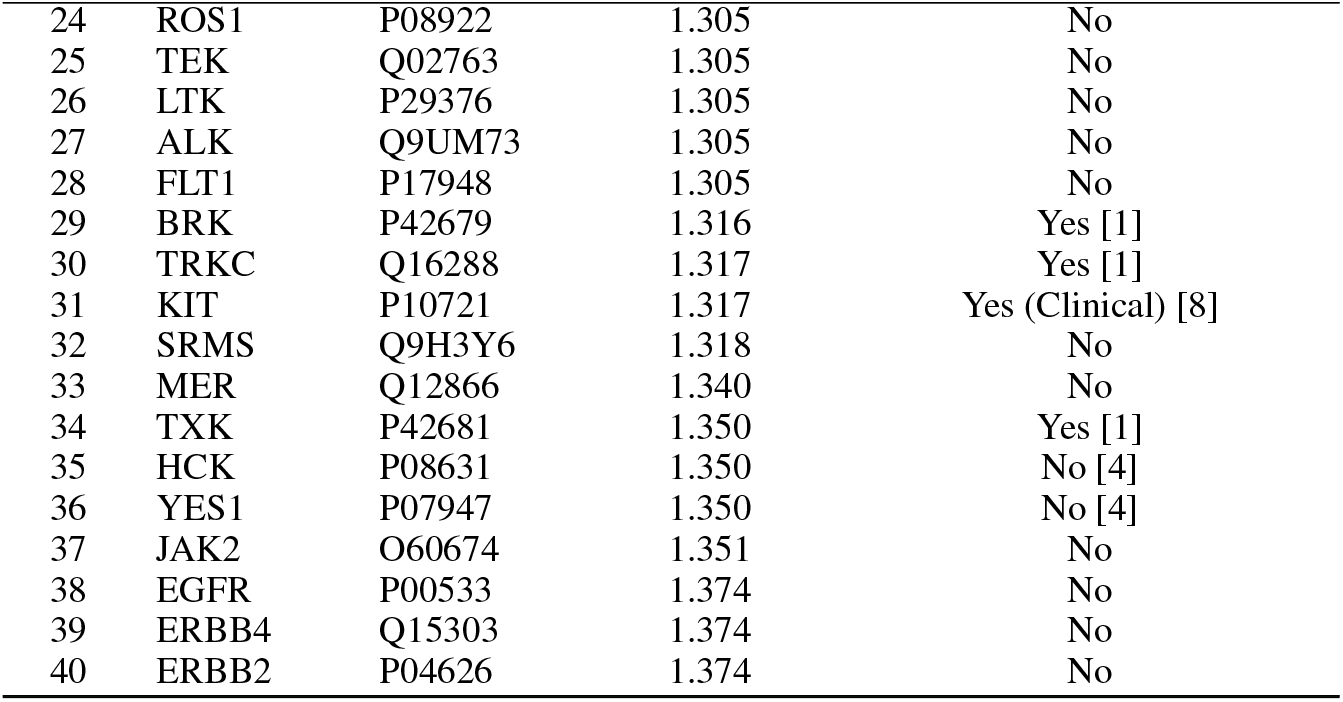
Top 40 structural hits for Imatinib-bound Abl kinase (1IEP) query against the Human Proteome. Ranking is based on LocAlign pLRMSD.

| Rank | Gene name | UniProt ID | pLRMSD (Å) | Known target? |
| --- | --- | --- | --- | --- |
| 1 | ZAP70 | P43403 | 0.000 | No (High structural similarity) |
| 2 | SYK | P43405 | 0.821 | No (High structural similarity) |
| 3 | ACK1 | P34925 | 0.964 | Yes [2] |
| 4 | ABL1 | P00519 | 1.096 | Yes (Primary Target) [3] |
| 5 | BLK | Q6J9G0 | 1.134 | Yes [1] |
| 6 | FRK | P42681 | 1.144 | Yes [1] |
| 7 | SRC | P12931 | 1.144 | No [4] |
| 8 | BMX | Q9H3Y6 | 1.167 | Yes [1] |
| 9 | CSK | P41240 | 1.167 | No |
| 10 | FGFR2 | Q16620 | 1.170 | Yes [2] |
| 11 | TRKA | P04629 | 1.170 | Yes [1] |
| 12 | FGR | P09334 | 1.170 | No [4] |
| 13 | DDR2 | Q16832 | 1.170 | Yes [6] |
| 14 | FGFR1 | P11362 | 1.170 | Yes [2] |
| 15 | FGFR3 | P22607 | 1.170 | Yes [2] |
| 16 | RET | P07949 | 1.193 | Yes [1] |
| 17 | FAK | Q05397 | 1.203 | No |
| 18 | FGFR4 | P22455 | 1.205 | No |
| 19 | LCK | P06239 | 1.230 | Yes [2] |
| 20 | MET | P08581 | 1.265 | No |
| 21 | PDGFRB | P09619 | 1.283 | Yes (Clinical) [7] |
| 22 | FGFR2* | Q16620 | 1.285 | Yes [2] |
| 23 | TIE1 | P35590 | 1.289 | No |
| 24 | ROS1 | P08922 | 1.305 | No |
| 25 | TEK | Q02763 | 1.305 | No |
| 26 | LTK | P29376 | 1.305 | No |
| 27 | ALK | Q9UM73 | 1.305 | No |
| 28 | FLT1 | P17948 | 1.305 | No |
| 29 | BRK | P42679 | 1.316 | Yes [1] |
| 30 | TRKC | Q16288 | 1.317 | Yes [1] |
| 31 | KIT | P10721 | 1.317 | Yes (Clinical) [8] |
| 32 | SRMS | Q9H3Y6 | 1.318 | No |
| 33 | MER | Q12866 | 1.340 | No |
| 34 | TXK | P42681 | 1.350 | Yes [1] |
| 35 | HCK | P08631 | 1.350 | No [4] |
| 36 | YES1 | P07947 | 1.350 | No [4] |
| 37 | JAK2 | O60674 | 1.351 | No |
| 38 | EGFR | P00533 | 1.374 | No |
| 39 | ERBB4 | Q15303 | 1.374 | No |
| 40 | ERBB2 | P04626 | 1.374 | No |

